# Dissecting polycomb complexes for enhanced fetal hemoglobin production

**DOI:** 10.64898/2026.04.16.718974

**Authors:** Paul J. Kaminski, Kristen Min, Elizabeth A. Traxler, Eugene Khandros, Osheiza Abdulmalik, Bailey Godfrey, Cheryl A. Keller, Belinda M. Giardine, Ross C. Hardison, Junwei Shi, Gerd A. Blobel

**Affiliations:** Division of Hematology, Children’s Hospital of Philadelphia, Philadelphia, PA; Division of Hematology and Oncology, University of Pennsylvania, Philadelphia, PA; Department of Biochemistry and Molecular Biology, Pennsylvania State University, University Park, State College, PA; Department of Cancer Biology, University of Pennsylvania, Philadelphia, PA

## Abstract

Polycomb repressive complexes PRC1 and PRC2 regulate diverse developmental processes, including the fetal-to-adult switch in hemoglobin production, a process whose reversal is a goal for the treatment of sickle cell disease and β-thalassemia. PRC inhibitors show promise for various disorders, but use is limited because of pleiotropic PRC activities. We explored whether fetal hemoglobin (HbF) can be reactivated in adult erythroid cells by selective perturbations of PRC1 or PRC2 components without complete loss of PRC function. A high-density CRISPR-Cas9 mutagenesis screen identified a region in the *EZH2* subunit where Cas9 induced exon 14 skipping (EZH2Δ14). EZH2Δ14, which lacks a portion of the CXC domain, relieves HbF repression while largely maintaining cellular fitness. EZH2Δ14 retains H3K27 methylation and repression of a PRC target gene subset. Experiments in cells derived from mice bearing human β-globin genes confirm that pathways mediating EZH2 control of HbF expression can function in a mouse model of *HBG* switching. These findings demonstrate that partial disruption of PRC can yield selective phenotypes, highlighting the therapeutic potential of targeting non-enzymatic domains within chromatin-modifying complexes.

**Key Points:**

- CRISPR-Cas9 screen across PRC1 and a saturating mutagenesis screen of PRC2 found the EZH2 CXC domain a desirable target for HbF induction
- the EZH2-CXC domain leads to exon 14 exclusion, resulting in de-repression of HbF but maintenance of cell fitness.

## Introduction

The fetal-to-adult hemoglobin switch occurs around birth when the fetal β-globin genes (*HBG1/2*) are silenced and the adult β-globin genes (*HBB* and *HBD*) are activated. Understanding this process is of clinical interest because the clinical outcomes of patients with sickle cell disease (SCD) and β-thalassemia can be greatly improved by replacing dysfunctional or inadequate levels of HBB containing hemoglobin (HbA) with HBG containing fetal hemoglobin (HbF)^1^. Initial genome wide association studies (GWAS) identified key regulators of the hemoglobin switch including the zinc-finger transcription factor *BCL11A*, a direct repressor of *HBG1/2* transcription that is the target for an FDA approved gene therapy for patients with SCD^2–4^. However, gene therapies are extremely resource intensive, and there remains a tremendous unmet need for safe and effective small-molecule HbF inducers.

While specific transcription factors including *BCL11A, ZBTB7A,* and *NFIX/NFIA* act directly at the β-globin locus, the role of epigenetic factors in globin switching is less well understood^5–7^. Histone modifying co-repressor complexes have been implicated in mediating *BCL11A* and *ZBTB7A* silencing at the β-globin locus, prominently including the nucleosome remodeling and deacetylase NuRD complex^8,9^. DNA methylation contributes to the recruitment and/or engagement of NuRD at the *HBG1/2* promoters, and targeted DNA demethylation is sufficient for *HBG* reactivation^10–13^. An siRNA screen linked multiple polycomb group proteins (PcG) to HbF repression^14^, while an independent CRISPR-Cas9 genetic screen identified the PcG gene *Bmi1* (PCGF4) as a novel HbF repressor^15^. PcG proteins assemble into distinct complexes: Polycomb repressive complex 2 (PRC2) contains EZH1 or EZH2, EED, and SUZ12 which methylate histone H3 at lysine 27 (H3K27)^16^. PRC1 binds to H3K27me3-decorated histones, ubiquitinates H2AK119 leading to chromatin compaction and gene repression^17^.

PcG proteins have critical roles in hematopoiesis. Loss of *Bmi1* leads to hematopoietic stem cell (HSC) exhaustion, while *Bmi1* overexpression enhances HSC expansion and erythroblast self-renewal^18–21^. PRC2 is critical for proper hematopoiesis during development, as *EED* has been shown to be necessary for HSC self-renewal in the adult bone marrow, while *EZH2* loss selectively impairs fetal hematopoiesis^22^. The switch from GATA2 to GATA1 in erythroid differentiation also coincides with a switch from EZH2 to EZH1 expression^23^.

Histone modifications differ between fetal and adult erythroblasts at developmentally regulated genes^24^. However, PcG proteins do not act directly on the β-globin locus, unlike the CpG rich α-globin cluster which is silenced by PcG proteins in non-erythroid cells^25,26^. Instead, PcG proteins modulate developmental expression of the β-globin genes indirectly. PRC2 and PRC1 co-operate to silence expression of the RNA binding proteins *IGF2BP1/3* and *LIN28B* in adult erythroid cells which in turn regulate BCL11A production ^15,27–29^. Clinical trials have begun targeting the EED subunit of PRC2 for HbF induction in patients with SCD. While targeting EED is moderately effective in inducing HbF in clinical trials^30^, inhibiting a core subunit of PRC2 shared among all PRC2 complexes may cause unwanted pleiotropic effects.

Our prior work utilized CRISPR-Cas9 screening technology to nominate novel HbF regulators and the pathways that they impinge upon^31–35^. Here, we hypothesized that specific domains within PcG proteins are selectively required for HbF repression in erythroid cells but dispensable for cell fitness. We designed a spCas9 based sgRNA library to comprehensively target nearly every protein domain-encoding region in PRC1, as well as nearly every PAM site in PRC2 core and accessory subunits. Our screens identified a critical region of *EZH2* that when targeted by Cas9 induced skipping of exon 14 which encodes a portion of the CXC domain of EZH2 (EZH2Δ14). EZH2Δ14 retains its catalytic SET domain, some level of methyltransferase activity, and chromatin association. Critically, EZH2Δ14 relieves HbF silencing while preserving repression of PcG target genes required for erythroid cell fitness. These findings highlight that selective perturbations of broadly acting chromatin modifying complexes can yield desirable phenotypic goals while reducing pleiotropic effects, opening a more focused therapeutic strategy to raise HbF expression in patients with β-hemoglobinopathies.

## Methods

### CRISPR-Cas9 sgRNA cloning and library construction

sgRNAs were designed with Benchling targeting coding regions of PcG genes. Libraries included six sgRNAs per PRC1 domain, saturating PAM mutagenesis of PRC2 genes, 100 non-targeting controls, and positive controls (BCL11A, ZBTB7A, ZNF410, MBD2). Oligos were cloned into pLRG2.1. Full lists are in Supplemental Tables.

### Cell culture

HUDEP2 cells were maintained and differentiated as described previously^36^. Expansion cultures were grown in StemSpan SFEM supplemented with SCF, EPO, doxycycline, dexamethasone, and penicillin/streptomycin. Differentiation was performed in a two-phase IMDM-based system with EPO, transferrin, insulin, and serum, with SCF withdrawal in phase 2.

Mobilized CD34⁺ HSPCs from healthy donors (Fred Hutchinson CCEH) were cultured using a four-phase erythroid differentiation protocol consisting of HSPC expansion, erythroid expansion, differentiation, and terminal maturation phases with staged cytokine adjustments.

### Lentivirus production and transduction

Lentivirus was generated in HEK293T cells by transient transfection of transfer plasmids with psPAX2 and VSV-G packaging constructs. Viral supernatant was collected at 72 hours and used to transduce HUDEP2 cells by spinoculation in the presence of polybrene. Transduced cells were sorted by flow cytometry based on fluorescent reporter expression.

### RNP electroporation of HSPCs

Cas9 RNP complexes were assembled in vitro and electroporated into primary CD34⁺ HSPCs using a 4D-Nucleofector system. Cells recovered in expansion media prior to erythroid differentiation. **CUT&RUN**

CUT&RUN was performed as described previously^37^ with minor modifications. Briefly, ConA-bound nuclei were incubated with primary antibodies, followed by pA-MNase digestion. Released DNA was purified, prepared into sequencing libraries, and sequenced (2×50 bp). Reads were aligned using bowtie2; peaks were called with SICER2 or MACS2, and differential binding assessed with DiffBind.

### RNA-seq and analysis

Stranded total RNA libraries were prepared following rRNA depletion and sequenced (2×50 bp, Illumina NextSeq). Reads were pseudo-aligned to hg38 using kallisto, and differential expression was performed using DESeq2 with ashr shrinkage. Genes with adjusted FDR < 0.05 were considered significant. Heatmaps were generated from normalized counts using z-score transformation. RNA-seq/CUT&RUN overlap analysis was performed using GeneOverlap.

### Flow cytometry

Cells were fixed, permeabilized, and stained with fluorescent antibodies to assess erythroid markers and HbF expression. Data were acquired on LSR-Fortessa or FACS-Canto instruments and analyzed in FlowJo.

### Artificial Intelligence Use

Claude Sonnet was used to assist with code debugging for NGS data analysis. Authors reviewed and take full responsibility for all content.

## Results

### PAM saturated CRISPR-Cas9 identifies the EZH2 CXC domain as a target for HbF induction

To uncover critical regions of PRC2 that cause increased HbF production while sparing cellular fitness, we generated a sgRNA library targeting nearly all PAM sequences in PRC2 core and accessory proteins, as well as most domains of PRC1-associated components and enriched cells based on HbF and persistence of sgRNAs in HUDEP2-Cas9 erythroid progenitor cells as described previously (**FIGURE 1a, SUPPLEMENTAL FIGURE1a-b**)^35^. MAGeCK^38^ normalized read counts from two biological replicate screens were used to generate parallel HbF enrichment and fitness scores for each sgRNA. HbF enrichment and fitness scores were calculated from log_2_fold changes in sgRNA abundance between sorted HbF-high vs. HbF-low cells, and between day-7 differentiated cells vs the starting sgRNA library (10 days expansion + 7 days differentiation) respectively. Using CRISPRO^39^, scores were aligned to the amino acid sequence of each protein. The most significantly enriched sgRNAs in HbF expressing cells targeted BMI1 and core PRC2 subunits EED/EZH2 (**FIGURE 1b-c**, **SUPPLEMENTAL FIGURE 1c-f).** Aside from BMI1, no other sgRNAs targeting a single PRC1 subunit induced high HbF levels. sgRNAs targeting accessory PRC2 subunits generally failed to induce high HbF levels, and most EZH2 targeting sgRNAs that triggered HbF enrichment also reduced fitness scores. However, we identified a region within the “CXC” domain of EZH2 in which multiple sgRNAs caused HbF enrichment but with near normal fitness scores (**FIGURE 1d, SUPPLEMENTAL FIGURE 1g)**. These sgRNAs clustered within amino acids 520-536 (hereafter referred to as sgEZH2-CXC), and compared to SET domain targeting sgRNAs showed limited fitness impairment but significant HbF enrichment over negative controls (**SUPPLEMENTAL FIGURE 8a-e).** Since all sgRNAs that scored in this region were within the same exon of EZH2, we hypothesized that Cas9-induced indels may cause exon exclusion, in this case exon 14. PCR analysis of cDNA using flanking exon primers or full-length *EZH2* cDNA sequencing of HUDEP2-Cas9 cells expressing sgRNAs targeting the EZH2-CXC domain revealed that exon 14 was selectively excluded in these cells (EZH2Δ14, **FIGURE 1e, SUPPLEMENTAL FIGURE 1h)**. We thus set out to characterize in depth the function of EZH2Δ14 in erythroid cells and assess the suitability of targeting the EZH2 CXC domain for HbF induction.

**Figure 1.**
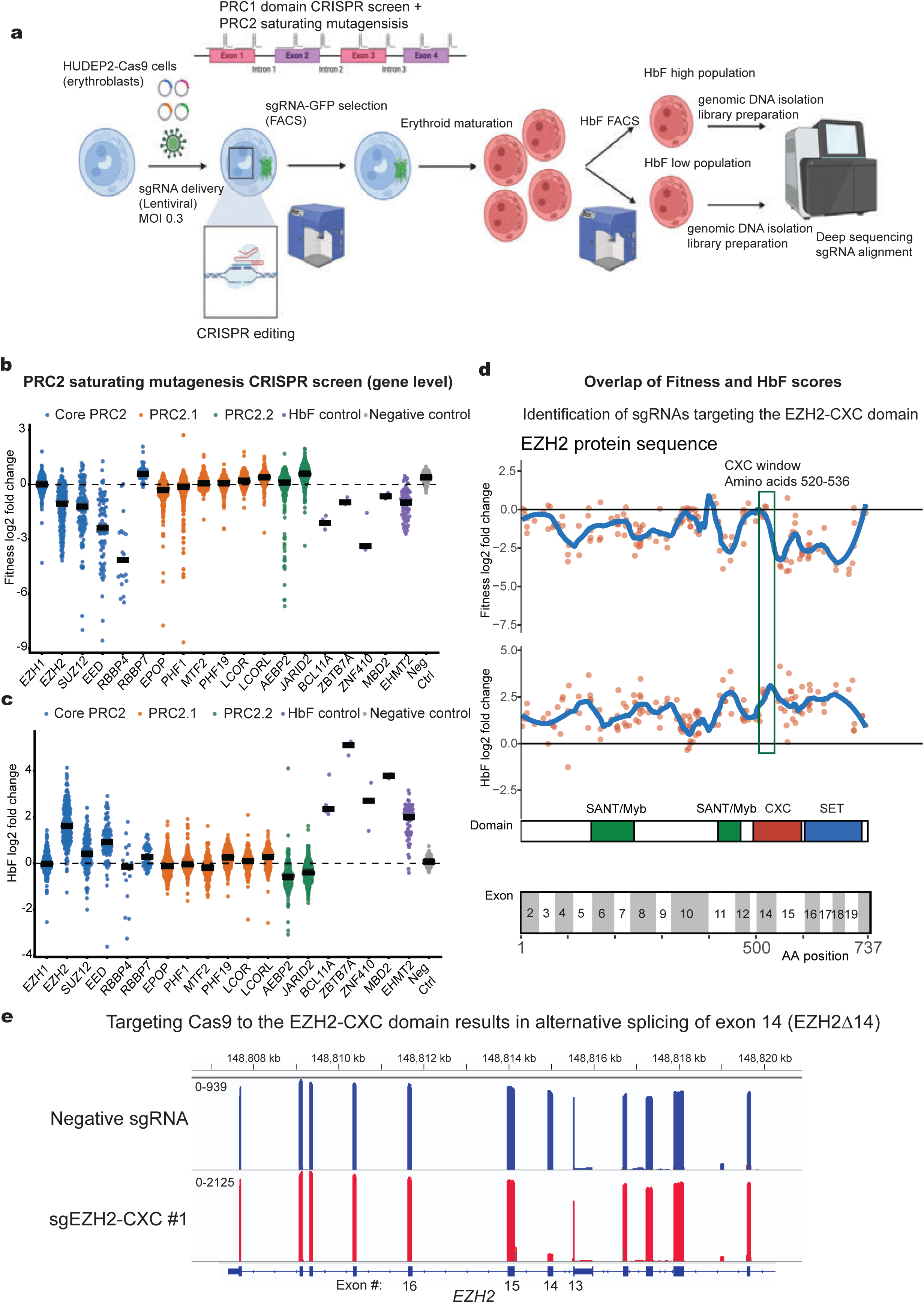
Saturating mutagenesis of the PRC2 complex uncovers the CXC domain of EZH2 as a specific HbF regulator with little effect on cell viability. **a)** Overview of CRISPR-Cas9 based screen in HUDEP2 cells. HUDEP2 cells stably expressing Cas9 were transduced n=2 biological replicates at low MOI (0.3) with LRG2.1-GFP sgRNA polycomb library. GFP positive cells were sorted and induced to differentiate for seven days prior to HbF FACS sorting and library deconvolution. Image created in BioRender **b-c**) Aggregated gene level scores for each sgRNA in the PRC2 saturating mutagenesis library for fitness scores (**b)** and HbF scores **(c)**. Horizontal bars indicate median score across all sgRNAs targeting a specific gene. **d)** HbF and Fitness scores for all sgRNAs targeting *EZH2.* Note a cluster of sgRNAs targeting the CXC domain of *EZH2* with high HbF enrichment scores and near zero fitness scores. **e)** Long read sequencing of EZH2 cDNA showing that sgRNAs targeting exon 14 of *EZH2* cause skipping of exon 14.

### Targeting the EZH2-CXC domain increases HbF expression in HUDEP2 cells

To validate the results from the CRISPR-Cas9 screen, we introduced into Cas9-expressing HUDEP2 three separate sgRNAs targeting the EZH2-CXC domain as well as three separate sgRNAs targeting the SET domain of EZH2, along with a positive control sgRNA targeting the BCL11A+58 enhancer and a non-targeting sgRNA as a negative control. Targeting the SET domain greatly destabilized EZH2, while targeting the CXC domain led to the production of EZH2Δ14 albeit at slightly lower levels (**FIGURE 2a)**. HUDEP2 cells targeted with sgEZH2-CXC retained slightly more H3K27me3 compared to sgEZH2-SET, but H3K27me3 was greatly reduced in both conditions. LIN28B, which is associated with increased HbF levels^15,28,29^, was elevated in both cases. Targeting either the CXC or SET domain of EZH2 resulted in comparable increases in HBG protein (**FIGURE 2b)**, in the number of cells producing measurable HbF (F-cells) (**FIGURE 2c-d, SUPPLEMENTAL FIGURE 2d),** and in HBG1/2 mRNA levels (**FIGURE 2e**).

**Figure 2.**
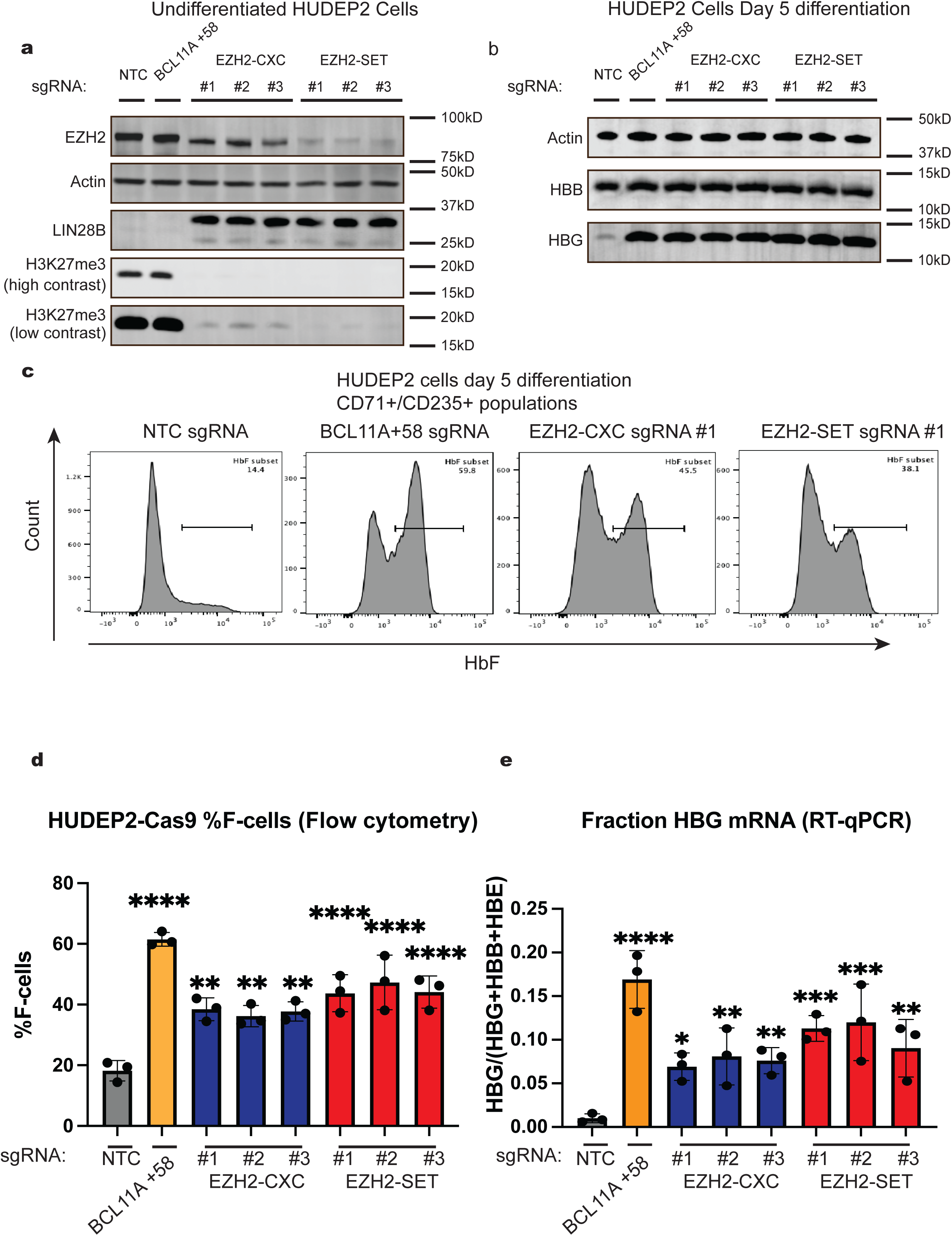
Targeted skipping of EZH2 exon 14 or loss of EZH2 induce fetal hemoglobin in HUDEP2 cells. **a)** Western blot with indicated antibodies of lysates from expansion phase HUDEP2-Cas9 cells targeted with three sgRNAs targeting the EZH2-CXC domain, three targeting the EZH2-SET domain, a positive control for HbF induction targeting BCL11A (BCL11A+58), or non-targeting (NTC) sgRNAs. **b)** Western blot with indicated antibodies of lysates from HUDEP2 cells transduced with indicated sgRNAs and differentiated for 5 days. **c)** Representative HbF flow cytometry of day 5 differentiated CD71+/CD235+ populations of HUDEP2 cells transduced with indicated sgRNAs. **d)** Aggregate data from n=3 biological replicates for % F-cells from flow cytometry experiments of day 5 differentiated HUDEP2 cells. Data shown are mean values with *p* values calculated by one way ANOVA followed by Fisher’s LSD test comparing to negative control. **e)** RT-qPCR analysis measuring the fraction of *HBG1/2* mature transcripts from n=3 biological replicates of day 5 differentiated HUDEP2 cells transduced with the indicated sgRNAs. Data shown are mean values with *p* values calculated by one way ANOVA followed by Fisher’s LSD test comparing to negative control.

We noticed that sgRNAs targeting the SET domain of EZH2 resulted in considerable spontaneous differentiation of HUDEP2 cells which was quantified by CD71/CD235 surface staining **(SUPPLEMENTAL FIGURE 2a)**. In contrast, cells expressing sgRNAs targeting the CXC domain of EZH2 exhibited significantly less spontaneous differentiation. Of note, targeting neither the EZH2 CXC nor the SET domain affected cell cycle profiles, suggesting that accelerated differentiation is not secondary to altered cell cycle progression (**SUPPLEMENTAL FIGURE 2b).** The Cas9 sgRNAs against the EZH2 CXC or SET domain did not target the paralog EZH1 **(SUPPLEMENTAL FIGURE 2c).** However, it is possible that the observed slightly elevated EZH1 levels are a reflection of EZH1/2 cross-regulation or compensation. We conclude that targeting the EZH2-CXC domain with Cas9 results in similar effects on HbF production, but with substantially milder effects on spontaneous differentiation of HUDEP2 cells.

### HUDEP2 cells expressing EZH2Δ14 retain H3K27me3 at a subset of PcG target genes

Targeting either the CXC or SET domain resulted in globally reduced H3K27me3 in HUDEP2 cells. However, the phenotypic difference between the two conditions suggests that EZH2Δ14 can place the H3K27me3 mark at a subset of relevant PcG genes. We performed CUT&RUN with αH3K27me3 antibody or IgG control in HUDEP2-Cas9 cells expressing negative control sgRNA, or sgRNAs targeting either the CXC or SET domain of EZH2. Because of global H3K27me3 changes, we included spike in *E. Coli* DNA^40^. As expected, targeting either the CXC or SET domain of EZH2 resulted in greatly reduced H3K27me3 genome wide (**FIGURE 3a**). To identify regions that retain H3K27me3 in the sgEZH2-CXC condition without arbitrary log-fold-change (LFC) or false discovery rate (FDR) cutoffs, we performed unsupervised k-means (k=3) clustering of peaks called from the negative control condition with deeptools using spike-in normalized bigwig tracks across all three conditions. While the majority of H3K27me3 peaks were reduced in both sgEZH2-CXC and sgEZH2-SET conditions, a cluster of peaks (cluster 1) retained H3K27me3 specifically in the sgEZH2-CXC condition (**FIGURE 3b)**.

**Figure 3.**
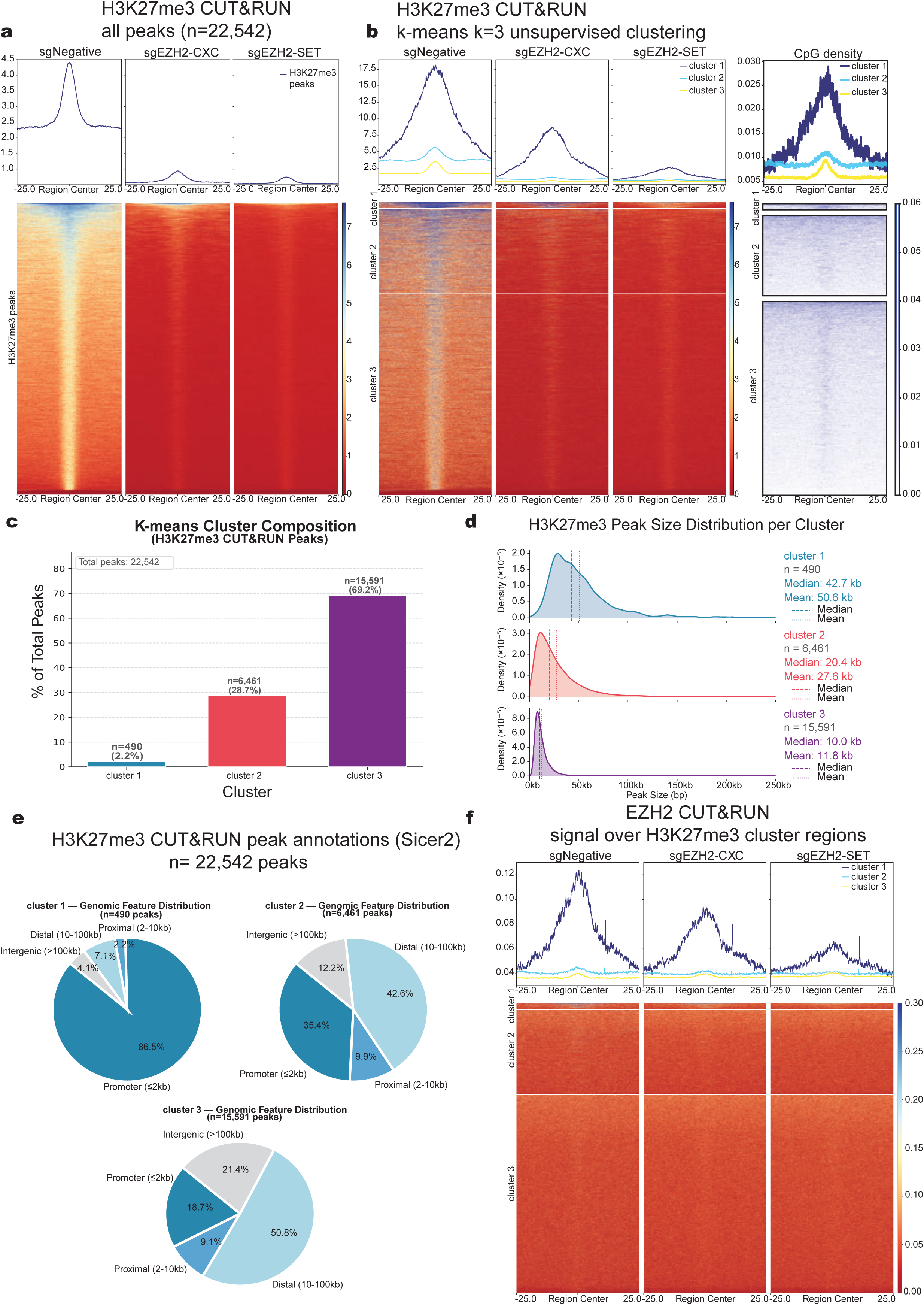
Retention of H3K27me3 at select PcG target genes in sgEZH2-CXC expressing HUDEP2 cells. **a)** CUT&RUN metaplots and heatmaps for HUDEP2 cells transduced with indicated sgRNAs displaying normalized signal intensity of H3K27me3 centered upon peaks called in the negative control condition. Plots are centered on the peak center and window sizes extend +/-25kb. BigWig heatmaps displayed are merged from n=3 biological replicates using E. Coli spike in derived normalization factors. **b)** CUT&RUN metaplots and heatmaps for HUDEP2 cells transduced with indicated sgRNAs displaying the normalized signal intensities for H3K27me3 in k=3 clusters using deeptools k means clustering. Plots are centered on the peak center and window sizes extend +/-25kb. BigWig heatmaps displayed are merged from n=3 biological replicates using E. Coli spike in derived normalization factors. **c)** Bar plot showing the total number of peaks and % of total peaks identified within each k means cluster. **d)** Density distribution of peak sizes within each individual k-means cluster. **e)** Classification of peaks within each k-means cluster based on relative distance to nearest transcription start sites (TSS) **f)** CUT&RUN metaplot and heatmaps for EZH2 normalized signal intensity in HUDEP2 cells transduced with the indicated sgRNAs centered on genomic regions from k means clusters identified in **b**. BigWig tracks are merged from n=3 biological replicates and are centered on peak center with window sizes +/-25kb

Unmethylated CpG islands present one mechanism by which PcG proteins are recruited to chromatin. Cluster 1 regions had higher CpG content compared to clusters 2/3. We additionally analyzed this dataset using DiffBind^41^ validating that the sgEZH2-CXC condition maintains H3K27me3 at specific regions **(SUPPLEMENTAL FIGURE 9a-c)**. We stratified peaks by cluster and found that cluster 1 represents a minority of peaks (490/22,542) that are wider (median 42.7kb) and more promoter proximal than clusters 2/3 (**FIGURE 3c-e).** Gene ontology analysis revealed distinct terms associated with clusters 1 and 2 (**SUPPLEMENTAL FIGURE 9d-f)**.

To test whether EZH2Δ14 occupancy correlates with the presence of retained H3K27me3, we performed anti-EZH2 CUT&RUN. This revealed that cluster 1 regions are bound by both EZH2Δ14 as well as the full-length form of EZH2 (**FIGURE 3f).** We orthogonally validated our CUT&RUN data by using ChIP-qPCR which confirmed retention of H3K27me3 in sgEZH2-CXC cells at the same genomic regions identified by CUT&RUN, such as at the *NR2F1*, *HOXA11,* and *ANKFY1* loci **(SUPPLEMENTAL FIGURE 3a-c)**. In contrast, the *JPH1* locus was confirmed to lose H3K27me3 in both knockout conditions (**SUPPLEMENTAL FIGURE 3d)**. Taken together, residual H3K27me3 at specific PcG target genes in EZH2Δ14 expressing HUDEP2 cells may explain the greater fitness compared to cells that are functionally null for EZH2.

### EZH2 exon 14 skipping increases HbF production in primary human erythroid cells

To test whether targeting the EZH2-CXC domain results in similar phenotypes in primary erythroid cells, we electroporated ribonucleoproteins into human primary CD34+ hematopoietic stem and progenitor cells (HSPCs) using Cas9 and sgRNAs targeting the CXC or SET domain of EZH2 as well as negative control and BCL11A+58 sgRNAs. sgRNA targeting of EZH2-CXC resulted in production of EZH2Δ14, while sgRNA targeting the SET domain of EZH2 resulted in near complete loss of EZH2 protein (**FIGURE 4a**). Importantly, in both conditions HBG protein levels were greatly increased in globin expressing late-stage erythroid cells (**FIGURE 4b).** The fraction of *HBG1/2* mRNA was elevated in both settings and reached levels similar to those observed in the BCL11A+58 sgRNA condition (**FIGURE 4c)**. We observed a two-to-three-fold increase in the fetal/adult hemoglobin (HbF/HbA) ratio in cells expressing sgEZH2-CXC or sgEZH2-SET (**FIGURE 4d)** and increased F-cell percentages (**FIGURE 4e, SUPPLEMENTAL FIGURE 4b**). Phenotypically, functional null mutations in *EZH2* resulted in accelerated differentiation of CD34+ cells compared to sgNegative control or sgEZH2-CXC. (**SUPPLEMENTAL FIGURE 4a).** sgEZH2-SET targeting impaired cell expansion during maturation stages while sgEZH2-CXC targeted cells proliferated largely normally (**SUPPLEMENTAL FIGURE 4c)**. We tracked the editing efficiency over time, which showed that indels within the SET domain were lost over the course of erythroid maturation while indels within the CXC region were stable, validating that cells lacking the EZH2-CXC domain do not undergo strong negative selection (**SUPPLEMENTAL FIGURE 10a-b)**. While an earlier study found loss of EZH2 causes defects in enucleation^42^, we observed that CD34+ cells edited at either the CXC or SET domain exhibited normal enucleation frequencies (**SUPPLEMENTAL FIGURE 10c).**

**Figure 4.**
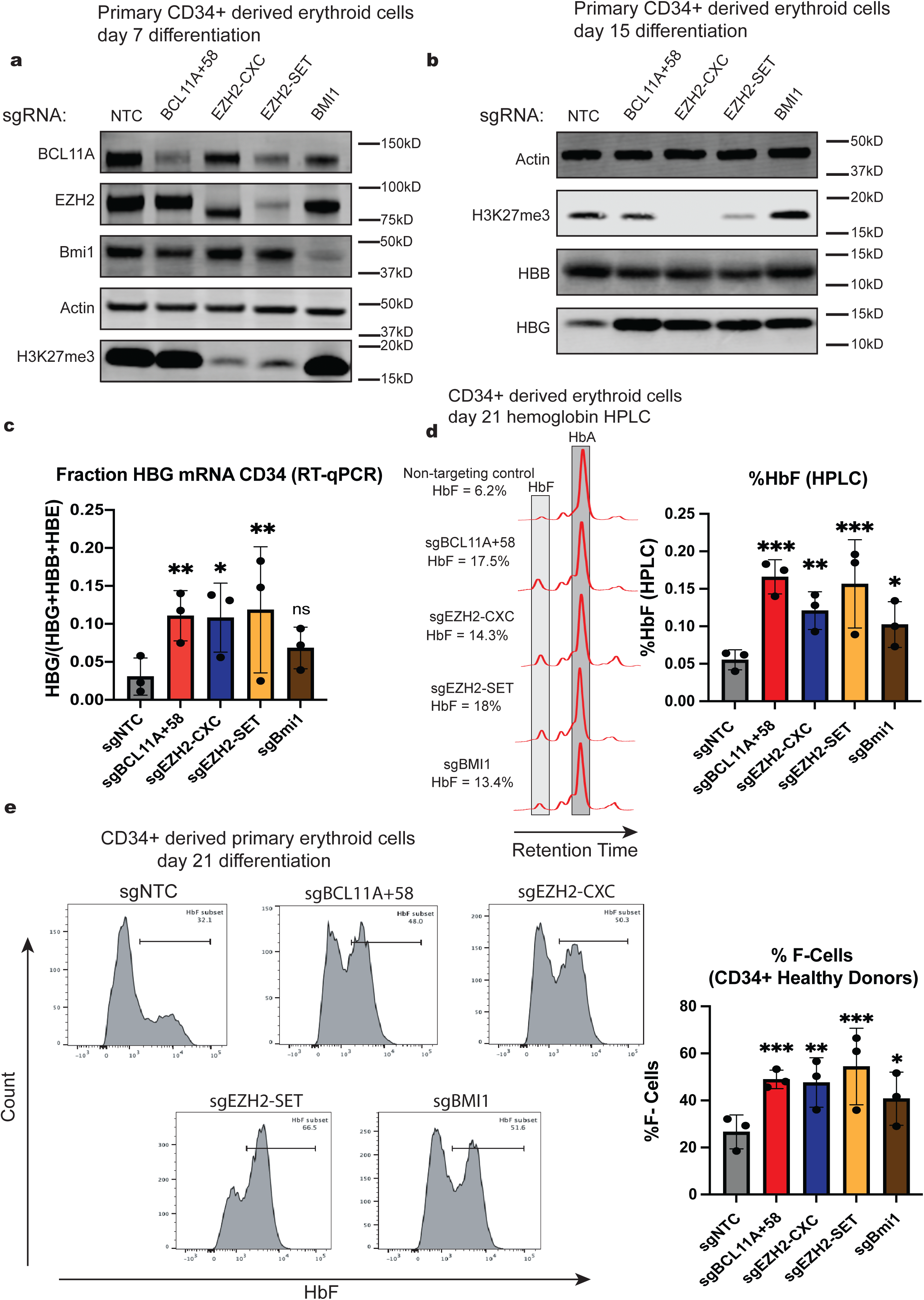
EZH2 exon 14 skipping in primary CD34+ derived erythroid cells increases fetal hemoglobin production. **a)** Western blot analysis of CD34+ cells at day 7 of erythroid differentiation following electroporation with the indicated sgRNAs. **b)** Western blot analysis of lysates from CD34+ cells at day 15 of erythroid differentiation following electroporation with the indicated sgRNAs. **c)** Fraction of *HBG1/2* transcripts in CD34+ derived erythroid cells at day 12 of differentiation following electroporation with the indicated sgRNAs. n=3 biological replicates. Data shown are mean values with *p* values calculated by one way repeated measures ANOVA followed by Fisher’s LSD test comparing to negative control. **d)** Representative hemoglobin HPLC traces from CD34+ derived erythroid cells at day 21 of erythroid differentiation following electroporation with the indicated sgRNAs. Aggregate HPLC analysis from n=3 biological replicates from CD34+ derived erythroid cells at day 21 of differentiation following electroporation with the indicated sgRNAs. Data shown are mean values with *p* values calculated by one way repeated measures ANOVA followed by Fisher’s LSD test comparing to negative control. **e)** Aggregate HbF flow cytometry analysis of CD34+ derived erythroid cells following 21 days of erythroid differentiation with the indicated sgRNAs. Data shown are HbF+ populations from CD71-/CD235+/Hoechst- from n=3 biological replicates. Data shown are mean values with *p* values calculated by one-way repeated measures ANOVA followed by Fisher’s LSD test comparing to negative control.

We compared CRISPR-Cas9 editing with current inhibitors of the PRC2 complex. Treatment with the EZH2 inhibitor EPZ-6438 or the EED inhibitor EED-226 at doses sufficient for HbF induction resulted in failure of in vitro erythroid expansion **(SUPPLEMENTAL FIGURE 4d-f)**. We conclude that pharmacological targeting of PRC2 mimics full genetic loss of EZH2 by increasing HbF while simultaneously disrupting erythroid cell maturation, whereas targeting the EZH2 CXC domain enables increased HbF production with less pronounced effects on primary erythroid cell viability and maturation.

### Fewer gene expression changes in sgEZH2-CXC compared to sgEZH2-SET

Since sgEZH2-CXC expressing HUDEP2-Cas9 cells had near normal fitness scores and retained H3K27me3 at select genomic regions, we hypothesized that this would be reflected in fewer changes in gene expression in sgEZH2-CXC expressing cells. We conducted bulk RNA-seq using HUDEP2-Cas9 cells expressing sgEZH2-CXC, sgEZH2-SET and control sgRNAs at day 5 of differentiation in three biological replicates. Using DESeq2 with an FDR<0.05, analysis of sgEZH2-SET vs sgNegative identified numerous gene expression changes compared to control (2095 differentially expressed genes – DEGs) with the majority (1309/2095 = 62%) of DEGs being upregulated **(FIGURE 5a, SUPPLEMENTAL TABLE 4).** In contrast, only 900 genes showed significant differences in expression for sgEZH2-CXC vs control, again with most DEGs being upregulated (640/900 = 71%) (**FIGURE 5b, SUPPLEMENTAL TABLE 5)**. Interestingly, 214 DEGs were detected between the sgEZH2-CXC and sgEZH2-SET condition, with 145/214 of the DEGs exhibiting higher expression in the latter (**FIGURE 5c, SUPPLEMENTAL TABLE 6**). Unsupervised hierarchical clustering of differentially expressed genes support this finding with many genes showing no significant change in expression in the sgEZH2-CXC condition when compared to the sgEZH2-SET condition (**FIGURE 5d**). We confirmed by RT-qPCR that three genes (*NR2F1*, *FGFR1* and the embryonic β-globin gene *HBE1*) were upregulated only in the sgEZH2-SET condition while maintaining repression in the sgEZH2-CXC condition (**SUPPLEMENTAL FIGURE 5a-c)**.

**Figure 5.**
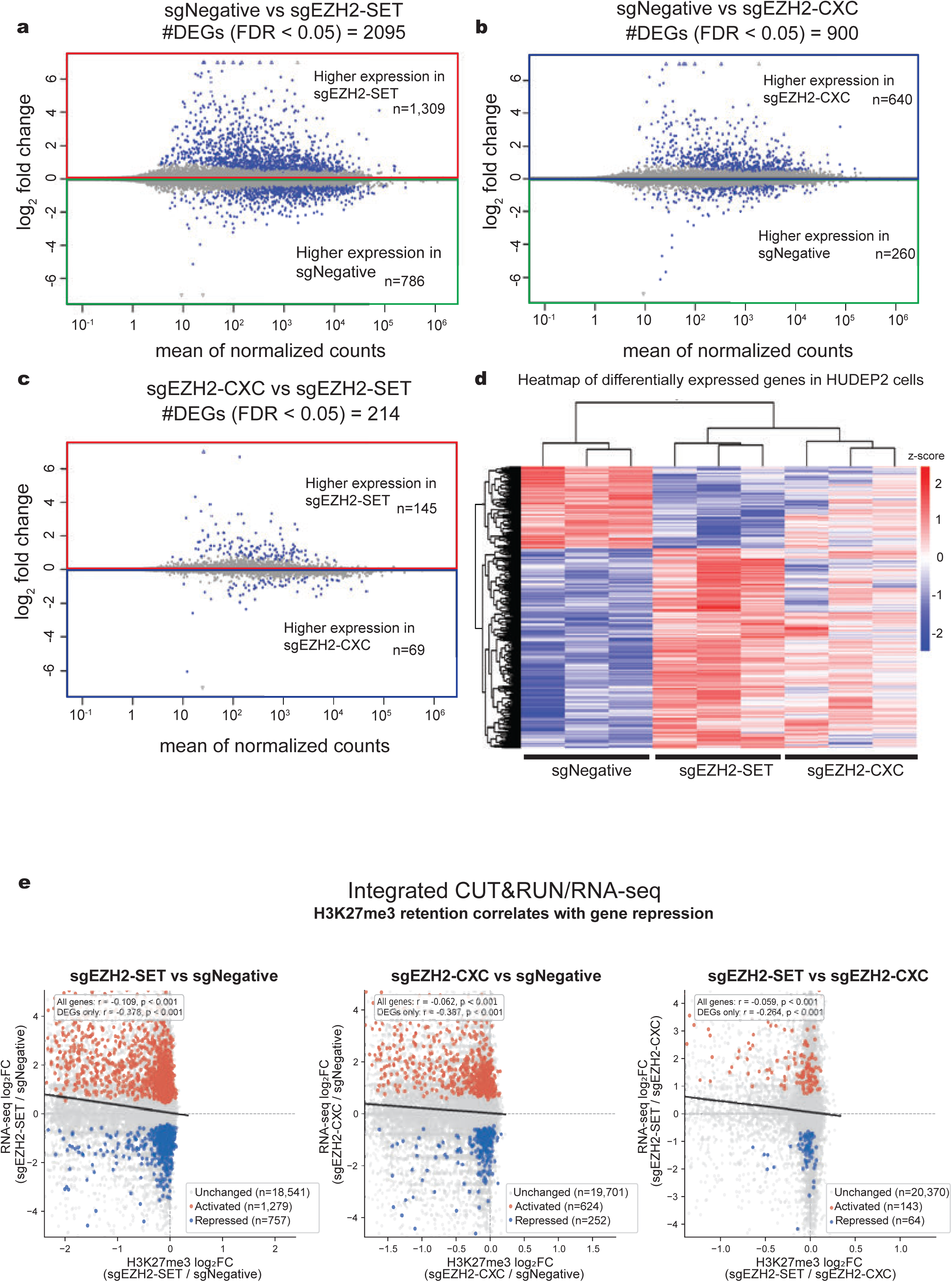
Reduced transcriptome changes upon EZH2 exon 14 skipping compared to full EZH2 loss. a-c) MA plots from RNA-seq experiments. Data shown are the DESeq2 normalized counts from. HUDEP2 cells at day 5 differentiation transduced with sgNegative, sgEZH2-CXC, and sgEZH2-SET sgRNAs. Points in blue are annotated as significant with FDR <0.05. The pairwise comparisons shown are sgNegative vs sgEZH2-SET (**a**); sgNegative vs sgEZH2-CXC (**b**); and sgEZH2-SET vs sgEZH2-CXC (**c**). RNA-seq experiments: n=3 biological replicates. d) Heatmaps and hierarchical clustering of differentially expressed genes from HUDEP2 cell RNA-seq. Normalized counts were used to generate visual heatmaps and clustering of data based on indicated sgRNA treatments and replicates. e) Comparison of H3K27me3 signal intensities (log_2_FC) at every annotated gene TSS +/- 3kb versus RNAseq log_2_FC for each annotated gene. Spearman correlations for the relationship between all genes are displayed as solid black lines. Highlighted genes are DEGs from RNAseq and are colored red when increased in expression, and blue if lower in expression in each comparison. Separate Spearman correlations for the DEG only subsets are also shown for each plot. Displayed p values are based on corresponding Spearman correlations and are calculated by a two-sided asymptotic t-test.

To examine the link between residual H3K27me3 and transcription, we plotted the change in H3K27me3 signal at all gene promoters against differential gene expression from RNAseq (**FIGURE 5e**). Loss of H3K27me3 was correlated with gene activation for all genes as well as the subsets of genes that are differentially expressed in each comparison. Importantly, in the sgEZH2-SET vs sgEZH2-CXC comparison, genes higher in expression in the sgEZH2-SET condition also exhibited greater loss of H3K27me3. We orthogonally used the GeneOverlap R package to assess overlapping gene sets between differential peaks from H3K27me3 CUT&RUN and bulk RNA-seq signal. Genes that remained repressed in the sgEZH2-CXC vs sgEZH2-SET condition maintained H3K27me3 (**SUPPLEMENTAL FIGURE 5d**). GO analysis of differential expressed genes from each comparison revealed unique terms for each comparison, but did not provide direct insight into which specific gene sets account for differences in cell fitness in sgEZH2-CXC targeted cells **(SUPPLEMENTAL FIGURE 5e-g).** Thus, targeting the EZH2 CXC domain resulted in substantially fewer transcriptional perturbations than SET domain disruption while still enabling de-repression of HbF, highlighting a mechanistic basis for selective epigenetic modulation.

### Ectopic EZH2Δ14 expression in EZH2 depleted cells restores erythroid maturation while preserving elevated HbF levels

Skipping exon 14 of EZH2 may affect EZH2 function but leaves open the possibility that lower levels of the EZH2Δ14 variant contribute to loss of H3K27me3 and gene expression changes. We therefore stably introduced into HUDEP2 cells vectors expressing Cas9-resistant forms of HA-tagged full-length EZH2 or EZH2Δ14, or empty vector. Subsequently, we introduced a sgRNA targeting the endogenous EZH2 SET domain or negative control sgRNA and measured H3K27me2 and H3K27me3 levels by western blot. Ectopic EZH2-WT fully restored global H3K27me2 and H3K27me3 (**FIGURE 6a-b**). In contrast, EZH2Δ14 expression at comparable levels failed to restore these marks (**FIGURE 6a-c**).

**Figure 6.**
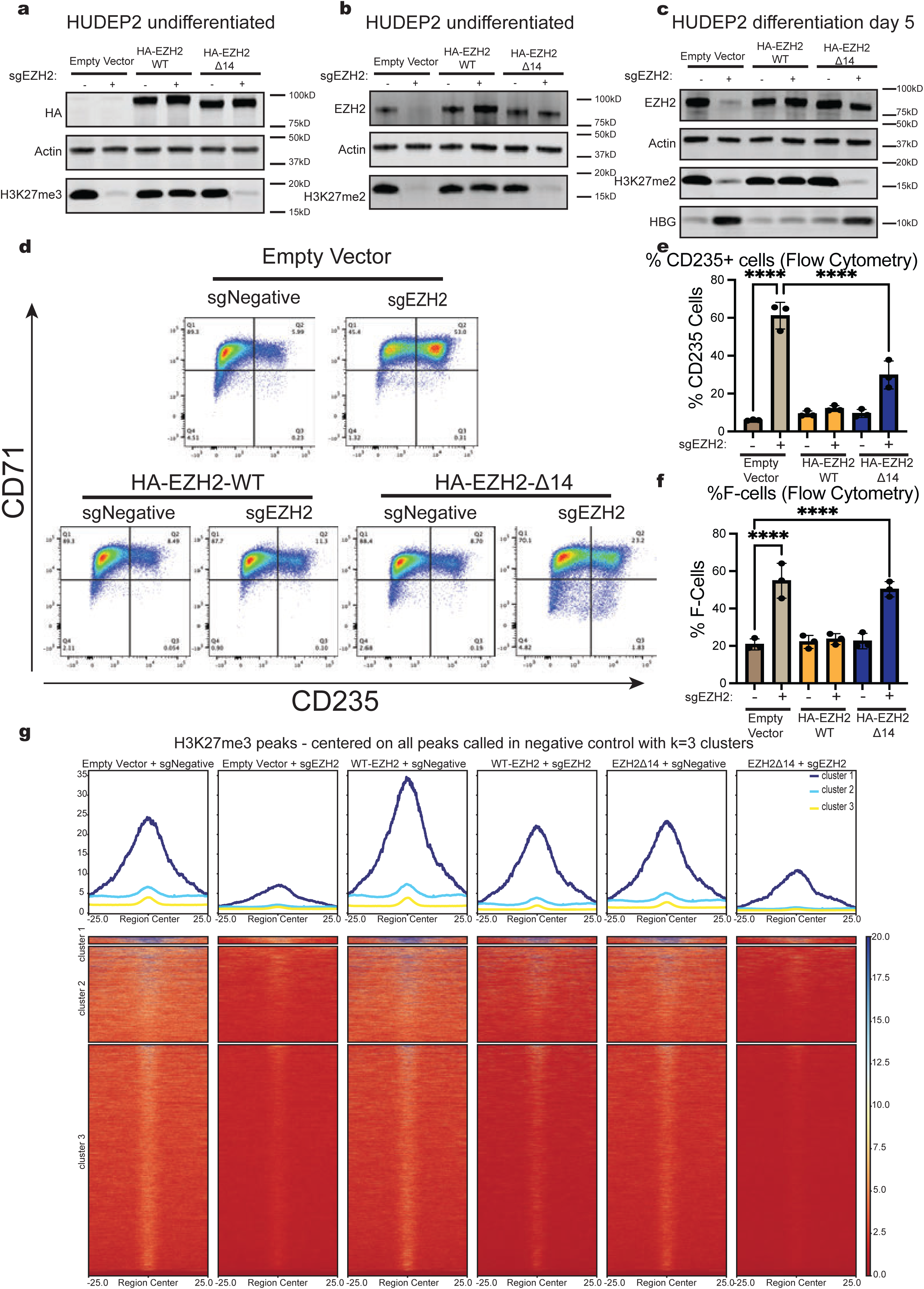
Ectopic EZH2Δ14 can partially compensate for EZH2 loss in HUDEP2 cells. **a-c)** Western blot of lysates from HUDEP2-Cas9 cells infected with empty vector or vector expressing HA-EZH2-WT, or HA-EZH2Δ14, transduced with indicated sgRNAs. HUDEP2 cells were analyzed in expansion phase (**a,b**) or after five days of erythroid differentiation (**c**). **d)** Representative flow CD71/CD235 cytometry plots of day 10 expansion phase HUDEP2-Cas9 cells infected with empty vector or vector expressing HA-EZH2-WT, or HA-EZH2Δ14, transduced with indicated sgRNAs. **e)** Aggregate data measuring spontaneous differentiation of EZH2 depleted HUDEP2-Cas9 cells infected with empty vector or vector expressing HA-EZH2-WT, or HA-EZH2Δ14. Percentages of CD235+ cells are shown. N=3 biological replicates. Data shown are mean values with *p* values calculated by one way ANOVA followed by Fisher’s LSD test. **f)** Aggregate data measuring the percentage of HbF+ in the EZH2 knockout-rescue HUDEP2 cells lines. N=3 biological replicates. Data shown are mean values with *p* values calculated by one way ANOVA followed by Fisher’s LSD test. **g)** CUT&RUN metaplots and heatmaps for H3K27me3 from EZH2 knockout-rescue HUDEP2-Cas9 cell lines. Data shown are H3K27me3 normalized intensities from n=3 merged biological replicates. Regions are centered on the peak center with +/-25kb window separated into k=3 clusters (same clusters as Figure 3b).

Importantly, EZH2-WT but not EZH2Δ14 reversed increased LIN28B and HbF synthesis as well as F-cell production (**FIGURE 6c,f; SUPPLEMENTAL FIGURE 6e,j)**. Moreover, ectopic EZH2-WT fully reversed the spontaneous differentiation as measured by CD71/CD235 surface staining (**FIGURE 6d-e)** whereas EZH2Δ14 reduced spontaneous differentiation by only 50% and failed to lower F-cell numbers (**FIGURE 6f**). We also tested whether EZH2Δ14 could re-establish repression of genes that were sensitive to exon 14 deletion in our RNA-seq dataset. RT-qPCR analysis showed that while EZH2-WT could rescue gene silencing of all tested genes equally (*HBG1/2, HBE, NR2F1, FGFR1*), EZH2Δ14 was unable to rescue the repression of *HBG1/2* but re-silenced *HBE1, NR2F1*, and *FGFR1* (**SUPPLEMENTAL FIGURE 6a-d**). We conclude that ectopic expression of EZH2Δ14 can partially uncouple HbF repression from defects in erythroid differentiation observed with functional loss of EZH2.

### Ectopic expression of EZH2Δ14 allows for retained methylation at specific H3K27me3 domains

To identify regions at which EZH2Δ14 can methylate H3K27me3, we performed anti-H3K27me3 CUT&RUN in EZH2 KO HUDEP2-Cas9 cells transduced with EZH2-WT, EZH2Δ14 or empty vector. We compared H3K27me3 enrichment genome wide across conditions using the same k=3 means clusters identified previously (**FIGURE 6g**). Loss of endogenous EZH2 dramatically reduced H3K27me3 genome wide which was rescued by ectopic WT-EZH2. Forced expression of EZH2Δ14 failed to globally establish H3K27me3 but was able to lay down this mark at a subset of regions (cluster 1). These regions overlap with those that retained H3K27me3 in the CRISPR-Cas9 experiments targeting sgEZH2-CXC, including the *SP9,* and *NR2F1* loci (**SUPPLEMENTAL FIGURE 6f-g**). Conversely, H3K27me3 was similarly lost at loci such as *JPH1* and *LIN28B* (**SUPPLEMENTAL FIGURE 6h-i**). Taken together, ectopically expressed EZH2Δ14 can maintain H3K27me3 at select polycomb target genes to maintain their repression.

### Preservation of PRC2 complex formation in the absence of EZH2 CXC domain

Deletion of protein domains may result in failure to form intact protein complexes. To test whether deletion of the CXC domain portion impairs assembly into the PRC2 complex, we performed co-immunoprecipitation experiments in HUDEP2 or HEK293T cells expressing 3x-FLAG tagged EZH2-WT or EZH2Δ14. This revealed that EZH2-WT and EZH2Δ14 were both capable of interacting with PRC2 core subunits EED, SUZ12, and RBBP4/7; and the PRC2 accessory subunits PHF19 and JARID2 (**SUPPLEMENTAL FIGURE 6k-m**). We conclude that EZH2Δ14 can be assembled into a functional PRC2 complex.

### PcG mediated control of *HBG* silencing is functional in a murine model system

To determine whether targeting the EZH2-CXC domain leads to similar phenotypes across species, we used a humanized mouse model where we crossed mice with homozygous human Townes configuration (HBA/HBB-HBG) with mice carrying homozygous insertions of Cas9^43,44^. We isolated the early erythroid progenitor cells from the fetal livers (FL) of compound heterozygous mice by lineage depletion of FL cells as described previously^45,46^. Negative control (sgRosa26), sgEZH2-CXC, sgEZH2-SET, or positive control (sgBCL11A) sgRNAs were introduced by retrovirus into lineage depleted FL cells which were expanded and differentiated for 48h (**SUPPLEMENTAL FIGURE 7a)**.

We verified successful editing of the mouse *EZH2* locus and found that sgEZH2-CXC targeted FL cells led to exon 14 exclusion (**SUPPLEMENTAL FIGURE 7e-f)**. FL cells transduced with sgRNAs targeting either the EZH2-CXC or SET domain expressed higher levels of human *HBG* transcripts by RT-qPCR (**SUPPLEMENTAL FIGURE 7b).** FL cells expressing sgRNAs targeting the *EZH2-*SET domain exhibited decreased cell expansion and accelerated maturation as measured by surface marker staining, while sgRNA targeting of the EZH2 CXC domain did not lead to significant effects on expansion or differentiation (**SUPPLEMENTAL FIGURE 7c-d)**. Taken together, these results indicate that pathways linking PcG to the regulation of the *HBG* genes are conserved in a mouse model of *HBG* switching.

## Discussion

Perturbing general chromatin modifying complexes for therapeutic goals is limited by pleiotropic functions. PcG complexes are being targeted for a variety of therapeutic goals, including the induction of HbF expression in patients with SCD^30^. Here we explored whether mutations or deletions within PcG proteins can relieve HbF repression in human adult erythroid cells, while preserving cellular fitness. Our approach was motivated by emerging evidence that specific subunits or domains within chromatin regulatory complexes can drive gene-selective phenotypes, supporting the concept that precision targeting of these elements may retain therapeutic efficacy while limiting off target effects^47–49^. We performed a saturating CRISPR-Cas9 based mutagenesis of all PRC2 core and accessory factor subunits and found that loss of the CXC domain of EZH2 markedly increased HbF production. The effects of CXC domain deletion on erythroid cell proliferation and differentiation were considerably milder than complete EZH2 loss. Deletion of the CXC domain was attributable to exclusion of exon 14, leading to the production of a shorter EZH2 variant (EZH2Δ14). A naturally occurring splice variant lacking this exon has been previously identified in male germ cells^50^, and in mouse brain^51^. In our hands in erythroid cells, EZH2Δ14 failed to globally restore either H3K27me2 or H3K27me3 in knock-out/rescue experiments. However, we did detect H3K27me3 at select regions in cells expressing EZH2Δ14, which was highly correlated with the maintenance of gene repression. Importantly, in EZH2 functional null cells these genes tended to lose H3K27me3 and were de-repressed.

Surprisingly, EZH2Δ14 expressing cells had markedly decreased H3K27me3 but only limited defects in erythroid maturation. Our findings are reminiscent of highly confined H3K27me3 patterns that have been found in tumors bearing H3K27M mutations or overexpressing the onco-histone mimetic EZH2 inhibitory peptide EZHIP. In these cases, most H3K27me3 is lost across the genome but is retained specifically at critical developmental promoters with CpG islands ^52–55^. PRC2 is initially recruited to such CpG island “nucleation sites” through the SUZ12 zinc-finger domain, where H3K27me3 is initially established before spreading to more distal regions^56–58^. Thus, EZH2Δ14 retains its ability to locate to the CpG islands but may fail to enable spreading of EZH2 mediated H3K27me3. Whether the spreading defect is due to decreased affinity for nucleosome substrates or allosteric activation of PRC2 will be an important question for future studies.

While global loss of PRC2 activity increases fetal hemoglobin levels in erythroid cells^14,59^, we found that full loss of EZH2 is associated with substantial defects in cell proliferation and differentiation in vitro. Our screening results suggest that high level HbF activation is only achievable upon loss of essential PRC2 subunits (including SUZ12, EED, and EZH2); however, targeting these subunits was generally poorly tolerated by HUDEP2 cells. A central finding of the present study is that the only sgRNAs that increased HbF while maintaining normal fitness scores were within the CXC domain of EZH2. These findings suggest that more selective targeting of PRC2 may reduce pleiotropic effects thus increasing the potential therapeutic window. While targeting of the EZH2-CXC domain has limited detrimental effects on the erythroid lineage, future studies will need to characterize the impact on the differentiation of other hematopoietic lineages. More broadly, our findings demonstrate that gene editing using Cas9 or base editors could be used to induce exon skipping in a variety of disease processes in which PcG proteins play a role. Interestingly, the sgEZH2-CXC sgRNA target sites encompass an exon splicing enhancer. This enhancer may be perturbed by antisense oligonucleotides (ASOs) to induce formation of EZH2Δ14^60^, which could be employed for more specific disruption of PRC2 functions. We envision that deeper interrogation of PcG complexes and advances in small molecule design may allow for more precise pharmacological targeting of PcG proteins for diverse pathological conditions, including β-hemoglobinopathies.

## Acknowledgements

This work was supported by NIH grants from the National Heart, Lung, and Blood Institute R01HL119479 to G.A.B., and F30HL178209 to P.J.K.; the National Institute of Diabetes and Digestive and Kidney Diseases R01DK058044 and R01DK054937 to G.A.B.; T32DK007780, and TL1DK143326 to P.J.K., and DK106829 to the Fred Hutchinson Cancer Research Center Cooperative. This project was additionally funded through the St. Jude Children’s Research Hospital Collaborative research consortium for novel gene therapies on sickle cell disease (CRC-SCD). We additionally thank the CHOP flow cytometry core for assistance with cell sorting experiments. HUDEP2 cells were gifted from R. Kurita and Y. Nakamura (RIKA BioResource Center).

We thank BioRender for generating the following figures used in the manuscript

https://BioRender.com/3812yhx

https://BioRender.com/guq6jwx

https://BioRender.com/psib1de

## Authorship Contributions

P.J.K., E.A.T., J.S., and G.A.B. conceived the study. P.J.K, K.M., E.A.T, and E.K. designed and performed experiments. P.J.K and K.M. collected and analyzed data. O.A. carried out hemoglobin HPLC experiments. E.K. and B.G. maintained mouse colonies and isolated murine fetal liver erythroid progenitor cells. C.A.K., B.M.G., and R.C.H. prepared RNA-seq libraries and sequenced all CUT&RUN libraries. R.C.H. and G.A.B acquired funding for the study. P.J.K. and G.A.B. wrote the manuscript with input from all authors.

## Data Sharing Statement

All raw and processed next generation sequencing data have been deposited into GEO. CUT&RUN data is available under accession number GSE316414 reviewer token: elsdcwcktpslzkx. RNAseq data is under accession number GSE316416 reviewer token: ujyvwscsnzafnsj.

