## Supplemental Materials for "Dissecting polycomb complexes for enhanced fetal hemoglobin production"

### Supplemental Methods

#### Mammalian cell culture

HUDEP2 cells were maintained and differentiated as described previously[1]. For the expansion phase of HUDEP2 cells, cells were grown at densities between 0.1 – 1.0 x 10^6 cells/mL in StemSpan serum free media (SFEM) with 50ng/mL supplemented human stem cell factor (SCF), 3U/mL erythropoietin (EPO), 1μg/mL doxycycline, 1μM dexamethasone, and 1% penicillin/streptomycin. HUDEP2 cells were induced to differentiate using a two-phase system. For phase 1, HUDEP2 cells were seeded at a density of 0.3 x 10^6 cells/mL and grown for 48 hours in Iscove's Modified Dulbecco's Medium (IMDM) based differentiation media with 3U/mL EPO, 5% fetal bovine serum, 320 μg/mL holo-transferrin, 2U/mL heparin, 10 μg/mL insulin, 1ug/mL doxycycline, and 50ng/mL human recombinant stem cell factor (SCF). For phase 2, cells were spun down and resuspended at 1.0 x 10^6 cells/mL in IMDM differentiation media without SCF for an additional 72 hours.

CD34+ human hematopoietic stem and progenitor cells (HSPCs) were mobilized and obtained from the peripheral blood of de-identified healthy volunteer donors from the Co-operative Center for Excellence in Hematology at the Fred Hutchinson Cancer Research Center. CD34+ cells were cultured in a four-phase culture system. CD34+ cells were thawed from cryopreservation and resuspended at density of 0.2 x 10^6 cells/mL in CD34+ Phase I media (HSPC expansion) using StemSpan SFEM supplemented with 100ng/mL SCF, 100 ng/mL thrombopoietin (TPO), 100ng/mL FLT-3L, 20ng/mL IL-6, 50ng/mL IL-3, 1x Glutamax, and 1% penicillin-streptomycin. HSPCs were grown in phase I for a maximum of five days and were then transitioned into phase II (erythroid expansion) containing StemSpan SFEM supplemented with 50ng/mL SCF, 3U/mL EPO, 10ng/mL IL-3, 40ng/mL IGF-1, 1μM Dexamethasone, 1x Glutamax, and 1% penicillin-streptomycin. Cells were allowed to expand for 10 days at densities between 0.1-1.0 x10^6 cells/mL in phase 2 before transition into phase 3 (erythroid differentiation) using IMDM-Glutamax base supplemented with 5% human AB serum, 10μg/mL insulin, 3U/mL heparin, 5U/mL EPO, 500μg/mL holotransferrin, 50ng/mL SCF, 1% penicillin-streptomycin, and 1μM Mifepristone. Cells were grown for four days in phase 3 media at densities between 0.5-5.0 x10^6 cells/mL and were then transitioned into phase 4 media (terminal erythroid differentiation) using IMDM-glutamax base supplemented with 5% human AB serum, 10μg/mL insulin, 3U/mL heparin, 5U/mL EPO, 500μg/mL holotransferrin, 1% penicillin-streptomycin, and 1μM mifepristone. Cells were maintained in phase 4 for five days to allow efficient hemoglobinization and enucleation before terminal analysis.

HEK293T cells were grown in DMEM supplemented with 10% FBS and 1% penicillin/streptomycin. Cells were maintained at 10% to 80% confluency prior to seeding for lentiviral production.

**Lentivirus production and infection of HUDEP2 cells**

sgRNAs cloned into pLRG2.1 (Addgene #108098) or EZH2 expression vectors (purchased from VectorBuilder) were used to produce lentivirus. Lentivirus was produced in HEK293T cells using a 3:2:1 ratio of psPAX2, VSVG, and transfer plasmid and a 3.5:1 ratio of PEI:DNA. Lentiviral particles were harvested 72 hours after transfection by collecting cell culture supernatant and spinning at 600g for five minutes to remove cellular debris. Lentiviral supernatant was then snap frozen or used directly to transduce HUDEP2 cells. To transduce HUDEP2 cells, 0.5 million HUDEP2 cells were incubated with 500uL of lentiviral supernatant, 8μg/mL polybrene, and 10uL HEPES in 1mL total volume of HUDEP expansion media. Incubated cells were then centrifuged in 24 well plates at 1,200g for 1.5 hours. Following transduction, cells were spun down and resuspended in fresh HUDEP2 expansion media. Transduced cells were then sorted for fluorescent marker expression 72 hours after transduction using a FACS Aria fusion sorter at the CHOP flow cytometry core.

**Flow cytometry**

Flow cytometry was used to assess erythroid maturation and measure HbF expression. 0.2-1 million HUDEP2 or primary erythroid cells were washed with PBS and then fixed with 0.05% glutaraldehyde. Cells were washed 3x with 0.2% BSA in PBS and then permeabilized with 0.1% Triton-X for 10 minutes on ice. Cells were washed 3x with 0.2% BSA prior to incubation with fluorescently tagged antibodies for 1hr at 4°C. Cells were washed 2x with 0.2% BSA in PBS prior to analytical flow analysis using FACS-Canto or LSR-Fortessa. FCS files were imported and processed using FlowJo.

**Cell cycle analysis**

1 x 10^6 HUDEP2 cells were collected by centrifugation and resuspended in 1mL room temperature PBS. Cell suspensions were added dropwise into 4mL of 100% ethanol on ice with vortexing. Cells were incubated in ethanol on ice for 10 minutes. Cells were pelleted by centrifugation, and washed with 5mL of room temperature PBS added to cells. After centrifugation, 50μL of 100μg/mL RNase was added to cells, and cells were stained with 200μL of 50μg/mL propidium iodide solution in PBS. Cells were analyzed for DNA content by flow cytometry and quantified in FlowJo.

**RT-qPCR for mRNA expression analysis**

RNA was isolated from 1-2 million HUDEP2 (day 5 differentiation) or primary CD34+ derived erythroid cells (day 13 differentiation) using the RNeasy mini kit. 100-250ng of purified RNA was used as template for cDNA synthesis using iScript master mix. cDNA was diluted 1:10 in nuclease free water for use in RT-qPCR. For analyzing relative gene expression, b-actin was used as internal reference control using the $2^{-\Delta\Delta CT}$ method, while for measuring globin mRNA abundance a relative globin proportion was calculated as HBG/(HBB+HBG+HBE) or HBE/(HBB+HBG+HBE) using ΔCT values. Primer sequences for RT-qPCR are listed in the supplemental tables.

**Western Blotting**

HUDEP2 cells grown during expansion phase or after 5 days of differentiation, and CD34+ cells at day 6 or day 13 of erythroid differentiation were used for western blot analysis. Cells were washed in 1x PBS and then lysed in RIPA buffer (25mM Tris-HCl pH 7.6, 150 mM NaCl, 1% NP-40, 1% sodium deoxycholate, 0.1% SDS) with supplemented 1X protease inhibitor cocktail (Sigma cat# P8340). Whole cell lysates were incubated on ice for 30min and then homogenized using a Bioruptor pico sonicator for 5 cycles. Lysates were then centrifuged at 15,000g for five minutes at 4°C, and the soluble protein lysate was removed and quantified using the Pierce BCA assay. Lysates were denatured with 1x LICOR buffer (LICOR cat# 928-40004) and incubated at 95°C for ten minutes. 5-10μg of protein was separated on 4-12% NuPAGE Bis-Tris gradient gels using 1x-MES-SDS buffer for 2hrs at 100V. Proteins were transferred onto PVDF membranes using 1X transfer buffer (25mM Tris, 192mM glycine, 10% methanol) for 90min at a constant 100V. Membranes were then stained using revert LICOR total protein stain (LICOR cat# 926-11011) , and the membranes were imaged for total protein to ensure consistent loading and total protein normalization. Stained membranes were reversed using 1x reversal solution (0.1M NaOH, 30% methanol). Membranes were blocked for one hour at room temperature on a shaker using 1x TBS blocking buffer (LICOR cat# 927-60001). After 3x washes in 1x TBST (1X TBS, 0.1% tween 20), membranes were incubated with primary antibody overnight at 4°C on a shaker in 1x TBST. The following day, membranes were washed 3x with TBST and incubated with fluorescent secondary antibodies for 1h at room temperature with shaking. After 3x washes with 1xTBST, membranes were allowed to dry before imaging on a LICOR Odyssey DLx. Antibody information is provided in supplemental tables.

**Ribonucleoprotein electroporation of primary HSPCs**

To induce CRISPR-Cas9 editing of primary HSPCs, 50 pmol of SpCas9 (IDT Alt-R SpCas9 cat#1081058) and 300 pmol modified sgRNA from Synthego were assembled in a PCR tube and incubated at room temperature for 15 minutes. 100,000-200,000 primary HSPCs at days 2-4 of expansion phase were washed in PBS, resuspended in P3 solution (Lonza P3 primary cell 4D-nucleofector, cat# V4XP-3032), and electroporated using a 4D Lonza nucleofector with program DZ-100. Following electroporation, CD34+ cells were allowed to recover in CD34+ expansion media for 24 hours prior to transition into erythroid expansion phase.

**Hemoglobin HPLC**

Erythroid cells at day 21 from CD34+ derived HSPCs were washed in 1X PBS and were lysed in nuclease free water at room temperature. Cell debris was removed by centrifugation at 15000g at 4°C for 10min and the soluble hemolysate was analyzed by high-performance liquid chromatography (HPLC). Hemolysates were separated on a weak cation-exchange column (Poly-CAT A: 50 mm x 4.6 mm, Poly LC, Columbia MD). Cleared lysates were analyzed with commercial standards consisting of equal amounts of HbF, HbA, HbS, and HbC (Helena Laboratories, Beaumont TX).

**RNA-seq library construction and data analysis**

RNA-sequencing libraries were prepared using the TruSeq stranded total RNA kit. 500ng of total RNA (RIN >8) were depleted for rRNA using NEB kit E7400. Purified RNA was fragmented, denatured, and reverse transcribed into cDNA followed by 8 cycles of PCR amplification. Barcoded libraries were pooled and sequenced at 2x50bp on the Illumina NextSeq 2000 platform. FastQ files from the sequencing run were pseudo-aligned and transcript level quantified to the hg38 cDNA reference genome using kallisto[2]. Differential expression between experimental conditions was performed in Rstudio using the DEseq2 package[3]. For visualization of differentially expressed genes (DEGs), we applied ashr log fold shrinkages to DEseq results[4]. To generate heatmaps of differentially expressed genes, we filtered for genes where padj<0.05 and |LFC| >1 in any comparison and calculated a $\log_2$(normalized count +1) for each gene across conditions. We then calculated z scores for each expression value and generated the heatmap using hierarchical row and column clustering with pheatmap. For categorical integration of RNAseq and CUT&RUN analysis, we used the GeneOverlap R package. Differentially expressed genes for each RNAseq pairwise comparison from DEseq2 were filtered for padj<0.05. Differential peaks from CUT&RUN Diffbind analysis were filtered for FDR<0.05 and were annotated using ChIPseeker[5], where we associated a peak with a gene if the peak center was within +/-3kb of the transcription start site of the gene. Filtered gene lists from both RNAseq and CUT&RUN were used to build contingency table and the odds ratio and p value of overlapping gene sets were calculated and displayed using drawheatmap.

**CUT&RUN DNA isolation, library preparation, and data analysis**

10μL Concanavalin beads (ConA) per sample were equilibrated in bead activation buffer (20mM HEPES-KOH pH 7.9, 10mM KCl, 1mM $CaCl_2$, 1mM $MnCl_2$) at room temperature. 0.5 million HUDEP2 cells were washed in 1X PBS and equilibrated in 100μL CUT&RUN wash buffer (20mM HEPES-NaOH pH 7.5, 150mM NaCl, 0.5mM Spermidine, 1X protease inhibitor) in 8-strip PCR tubes. Cells were mixed with ConA beads and allowed to bind to beads for 10min at room temperature. The ConA-cell mixture was magnetically separated and washed 3X in wash buffer, and the bead-cell mixture was resuspended in 50uL cold antibody buffer (1x CUT&RUN wash buffer + 0.08% digitonin +2mM EDTA). Respective antibodies were added and samples were incubated overnight at 4°C on a rotating nutator. The next day, samples were placed on a magnetic separator and washed 3X with 200μL cold CUT&RUN digitonin buffer (1x CUT&RUN wash buffer + 0.08% digitonin). Cell/bead mixtures were then resuspended in 50μL of cold digitonin buffer and incubated with 2.5μL of 20x CUTANA p-AG-MNase (Epicypher SKU: 15-1016) for 10min at room temperature. Cell/bead mixtures were washed 3x using digitonin wash buffer and then resuspended in 50μL of digitonin wash buffer and moved into a cold room on ice. MNase cleavage was initiated by adding 1μL of cold 100mM $CaCl_2$ (2mM final concentration) to each sample. MNase digestion was allowed to proceed for 90min before addition of 33μL of STOP buffer (340mM NaCl, 20mM EDTA, 4mM EGTA, 50μg/mL RNase A, 25 μg/mL glycogen) with 0.25ng E. Coli spike in DNA (Epicypher SKU: 18-1-401) per reaction. Cleaved DNA was released by incubation at 37°C for 10min, and cleaved DNA was purified from the soluble fraction using Qiagen QIAquick PCR clean up (Qiagen cat#28104). Purified DNA was quantified by Qubit high sensitivity reagents and ~10ng was used for library construction using barcoded KAPA hyperprep kits (KK8504). Pooled libraries were sequenced on 2x50bp Illumina NextSeq platform. FastQ files were aligned to the hg38 reference genome (or E. coli reference for spike in) using bowtie2[6]. Alignment sam files were then converted to bam format using samtools, and duplicate reads were removed using picard. De-duplicated bam files were visualized using deeptools bamCoverage. For bigWig normalization of CUT&RUN tracks, the normalization factor was determined using the relative reference reads per million method (RRPM) of exogenous spike in E. Coli reads[7]. Peaks were called within individual biological replicates using MACS2 (broadpeak) or SICER2 for experimental samples with IgG CUT&RUN samples used as background[8]. For heatmap and metaplot visualization, scaled bigWig tracks were plotted using peaks called with SICER2 in the merged negative control condition. For differential binding analysis, replicate bam files were analyzed in DiffBind[9] using default settings with a consensus peak set called using MACS2.

**Integration of CUT&RUN and RNAseq data**

To compare the change in signal for H3K27me3 across all genes, transcription start sites for each gene were extracted and a window +/- 2kb were used to create a bed file to define promoter regions. The signal intensity from individual normalized bigwig files were calculated using multiBigwigSummary. The mean value of each bigwig within the defined promoter region was used to calculate a $\log_2$fold change for each pairwise comparison. RNAseq $\log_2$fold change data from DEseq2 outputs was then merged with the CUT&RUN $\log_2$ fold change (LFC) to generate a combined CUT&RUN LFC and RNAseq LFC for each gene. Combined scatter plots were generated for each pairwise comparison plotting the H3K27me3 $\log_2$FC and RNAseq LFC. Spearman correlations and two sided p-values were calculated based on every gene, as well as for genes that had been subset as differentially expressed genes (DEGs) with padj<0.05 from RNAseq.

**Gene Ontology analysis**

For RNAseq datasets, Gene Ontology (GO) enrichment of differentially expressed genes (DEGs) was performed using clusterProfiler (v4) in R with org.Hs.eg.db as human annotation. Genes were filtered based on adjusted p values <0.05 and LFC >1.0. Enrichment was tested specifically for upregulated genes using enrichGO with Benjamini-Hochberg correction. All ontology categories were tested. The background universe was defined as all expressed genes with a non-NA padj value. Redundant GO terms were filtered out and results were visualized with bar plots and enrichment map plots.

For CUT&RUN datasets, GO enrichment of H3K27me3 k-means clusters was performed using GREAT (Genomic Regions Enrichment of Annotations Tool)[10] v4.0.4. Peaks from each cluster were submitted individually to GREAT using hg38 genome assembly and default association rule. The complete Sicer2 H3K27me3 peakset (n=22,542 peaks) was used as custom background. Significant enrichment was defined as binomial FDR q-value <0.05 for GO biological process terms.

**Chromatin Immunoprecipitation, quantitative PCR (ChIP-qPCR), and ChIP-seq**

5-10 million HUDEP2 cells grown in expansion phase were washed 1x in PBS and were crosslinked in 1% formaldehyde in PBS for 10 mins at room temperature. Crosslinking was quenched by addition of glycine to final concentration of 1M. Cells were lysed in ChIP lysis buffer (10mM Tris pH 8.0, 10mM NaCl, 0.2% NP-40, 1x protease inhibitors) and nuclei were enriched by centrifugation at 10,000g for 2 minutes. The supernatant was removed and nuclei lysed using nuclear lysis buffer (50mM Tris pH 8.0, 10mM EDTA, 1% SDS, 1x protease inhibitor) for 20 minutes on ice. Nuclear lysate was then transferred to sonication tubes and chromatin was sheared using 5x cycles (30 seconds on, 30 seconds off) using the bioruptor pico sonicator. Sonicated chromatin was centrifugated at 10,000g for 5 minutes at 4°C to pellet cellular debris. Soluble chromatin was transferred to 15mL conical tubes and diluted to 4mL in IP dilution buffer (20mM Tris pH 8.0, 2mM EDTA, 150mM NaCl, 1% Triton-X, 0.01% SDS, 1x protease inhibitor). Diluted chromatin was pre-cleared using 50μL of protein A/G beads and 50μg of isotype IgG control for 2 hours at 4°C. Pre-cleared chromatin was then enriched by centrifugation, and 900μL of chromatin was used for each immunoprecipitation with 5μg of specific antibody and 35μL of protein A beads. Chromatin immunoprecipitation occurred overnight at 4°C on a rotator. The following day, ChIP reactions were separated using a magnetic separator and washed once with IP wash buffer (20mM Tris pH 8.0, 2mM EDTA, 50mM NaCl, 1% Triton-X, 0.1% SDS), followed by 2x washes with high salt wash buffer (20mM Tris pH 8.0, 2mM EDTA, 500mM NaCl, 1% Triton X, 0.01% SDS), 1x wash with IP wash buffer #2 (10mM Tris pH 8.0, 1mM EDTA, 0.25 M LiCl, 1% NP-40, 1% sodium deoxycholate). ChIP samples were then equilibrated with 2x washes in TE buffer, and then eluted in a final volume of 200 μL of ChIP elution buffer (100mM $NaHCO_3$, 1% SDS). Input samples and ChIP samples were then treated with RNase A (0.1 mg/mL) and de-crosslinked for 3h at 65°C. Proteinase K was then added to final concentration 0.3mg/mL and de-crosslinked for an additional 1hr at 65°C. DNA was then purified using QIAGEN QIAquick PCR cleanup kits and quantified by Qubit. Samples were diluted 1:5 in nuclease free water and analyzed by qPCR using region specific primers. Enrichment was calculated using the percent input method. For library preparation, 5ng of ChIP DNA was used for input into KAPA hyperprep kits (KK8504) or NEB next library prep kits (E7645). Libraries were then pooled and sequenced 2x50bp on the Illumina NextSeq platform.

**Co-immunoprecipitation**

Whole cell lysates were prepared from expansion phase grown HUDEP2 cells lysed in cold IP lysis buffer (25mM Tris-Cl, pH 7.5, 150mM NaCl, 1mM EDTA, 1% NP-40, 5% glycerol, 1x Protease inhibitor Sigma cat# P8340). Whole cell lysate was incubated for 30min on ice and the soluble whole cell lysate was quantified by BCA. 5% of input (approximately 25μg) of lysate was saved for input controls. For each immunoprecipitation reaction, 50μL of Protein-A conjugated magnetic beads (ThermoFisher Dynabeads cat#10001D) were transferred to sterile 1.5mL Eppendorf tubes. Beads were equilibrated using 500μL of cold IP lysis buffer. Beads were magnetically cleared of supernatant and 5μg of antibody was added to beads in a volume of 200μL of cold lysis buffer. Antibody-bead mixtures were incubated at room temperature for 2 hours on a spinning rotator. Mixtures were then magnetically separated and washed using 200μL cold IP wash buffer (25mM Tris-Cl pH 7.5, 150mM NaCl, 1mM EDTA). The wash buffer was removed and 500μg of prepared whole cell lysate was incubated with antibody bead mixtures overnight at 4°C on a rotator. The following day, the mixture was magnetically separated, and bead mixtures were washed 3x using 200μL cold wash buffer. To competitively elute captured 3x-FLAG tagged proteins, 100μg of purified 3x-FLAG peptide (Sigma cat#F4799) was added in a volume of 20μL of Tris-buffered Saline (25mM Tris-Cl pH 7.5, 150mM NaCl). Elution reactions were incubated on ice for 2 hours, and soluble fractions were magnetically separated and collected for downstream western blot analysis.

**EZH2 Alternative splicing analysis**

To measure alternative splicing of EZH2, total RNA was purified from 2M HUDEP2 cells using Qiagen RNAeasy kits. cDNA was generated using 150-250ng of purified RNA using iScript master mix. Full length cDNA of EZH2 was generated by PCR amplification of EZH2 using primers on the 5' and 3' end of EZH2 with phusion flash polymerase. PCR products were run on a 1% agarose gel and the full length EZH2 cDNA PCR products were purified with Qiagen Gel extraction kits. Long read nanopore sequencing of the full-length products was performed by Eurofins genomics. To align sequencing reads, the fastq files were aligned to the human reference cDNA index using minimap2[11]. Aligned reads were visualized on IGV to assess read coverage and splicing frequency.

PCR based measurement of *EZH2* splicing was performed using cDNA from HUDEP2 cells with primers spanning the flanking exons (exons 13 and 15) using Phusion flash polymerase. PCR products were visualized on a 2% agarose gel, and the relative splicing frequency was inferred. Primers used for PCR analysis are listed in supplementary tables.

**Mouse fetal liver isolation**

Mouse strains were obtained from Jackson Labs: Townes strain 027265, $\beta^A$, Rosa26-Cas9 strain 026179). Erythroid progenitor cells were isolated from E14.5 fetal livers of Townes/Cas9 crosses by antibody mediated lineage depletion as described previously[12,13]. Isolated fetal liver cells were immediately transduced with pSL21 sgRNA retroviral carrying a VEX-GFP reporter. 48-96 hours following transduction, VEX-GFP positive cells were sorted by FACS and placed in fetal liver expansion media for 8-10 days. Fetal liver cells were induced to differentiate for 48h prior to downstream analysis. All mouse experiments were conducted in compliance with IACUC protocols at the Children's Hospital of Philadelphia.

**Inference of CRISPR editing (ICE analysis) for indel efficiency**

To measure the relative frequency of indels in CRISPR-Cas9 experiments, genomic DNA from edited cells were harvested using Invitrogen PureLink Genomic DNA purification kits (cat# K182002). 10ng of purified genomic DNA was then used as input for PCR reactions using primers designed specifically to amplify starting from intronic regions upstream and downstream of the target cut site. Negative control PCR reaction in unedited cells were used as a control reaction and both PCR tracers (.ab1 files) were uploaded to the ICE[14] analysis portal to assess editing efficiency.

**Statistical analysis**

Statistical analysis of numeric data was performed using the indicated statistical tests with graph-pad Prism. P values shown on figures are: * $p<0.05$, ** $p<0.005$, *** $p<0.0005$, **** $p<0.00005$.

**SUPPLEMENTAL FIGURE 1**

**a)** Flow cytometry gating strategy for the CRISPR-Cas9 screen in HUDEP2 cells. Viable cells are gated by FSC-A/SSC-A, singlets by FSC-A/FSC-H, followed by CD235+(PE-Cy7), CD71+(PE-A)/FSC-Low, and HbF-Hi/Low(APC-A)

**b)** Principal component analysis of normalized read counts for the sgRNA sequencing libraries before viral packaging (plasmid), immediately after sgRNA library transduction (undiff), after five days of expansion and 7 days of erythroid differentiation (diff7), and HbF Hi/Lo.

**c-d)** MAGeCK gene level rankings for sgRNAs enriched in the HbF Low (**C**) and HbF High (**D**) sequencing libraries

**e-f)** Gene level aggregated fitness scores (**e**) and HbF scores (**f**) for each sgRNA in the PRC1 domain targeted CRISPR-Cas9 screen. Horizontal bars represent the median value for all sgRNAs targeting a specific gene.

**g)** Plot of EZH2 fitness and HbF enrichment scores of every sgRNA targeting EZH2. Each point represents a particular sgRNA, with a LOESS regression shown in a blue line with a 99% Confidence interval in gray. sgRNAs are colored by annotated domain and labeled by the predicted Cas9 cleavage codon.

**h)** PCR based analysis of *EZH2* alternative splicing in HUDEP2 cells following CRISPR-Cas9 editing with the indicated sgRNAs

**SUPPLEMENTAL FIGURE 2**

**a)** Flow cytometry plot of HUDEP2-Cas9 cells transduced with respective sgRNAs analyzed 12 days post transduction to monitor spontaneous differentiation vis CD71/CD235 surface staining

**b)** Propidium iodide staining of HUDEP2-Cas9 cells transduced with respective sgRNAs to assess cell cycle state of HUDEP2 cells.

**c)** Immunoblots with indicated antibodies of protein lysates from HUDEP2 cells edited with indicated sgRNAs**.**

**d**) Median fluorescence intensity (MFI) of HbF-APC measured by flow cytometry in the indicated cell lines. Bars represent mean values +/- standard deviation.

**SUPPLEMENTAL FIGURE 3**

**a-d)** ChIP-qPCR validation of retained and lost peaks from analyzed CUT&RUN datasets. CUT&RUN tracks are visualized for the *NR2F1* (**a**), *HOXA11* (**b**), *ANKFY1* (**c**), and JPH1 (**d**). ChIP qPCR was performed using the amplicons labeled for n=3 biological replicates and input normalized to assess congruence between CUT&RUN and ChIP datasets. Labeled plots on right hand side are the percent input for each condition with mean value +/- standard deviation.

**SUPPLEMENTAL FIGURE 4**

**a)** Representative flow cytometry plots to assess CD71/CD235 surface marker staining of primary CD34+ cells from healthy donors on day 9 of erythroid differentiation following RNP electroporation with Cas9 and respective sgRNAs.

**b**) Median fluorescence intensity (MFI) of HbF-APC measured by flow cytometry in the indicated cell lines. Bars represent mean values +/- standard deviation.

**c)** Aggregate cell count data from n=3 healthy CD34+ donors displaying erythroid cell expansion following RNP electroporation with respective sgRNAs

**d)** Aggregate cell count data from n=2 healthy CD34+ donors displaying erythroid expansion with continual treatment of cells with commercially available PRC2 inhibitors.

**e)** Representative hemoglobin HPLC traces from CD34+ cells treated with respective concentrations of commercially available PRC2 inhibitors.

**f)** Western blot analysis of CD34+ cells treated with respective concentrations of PRC2 inhibitors after 15 days of continuous drug treatment during differentiation.

**SUPPLEMENTAL FIGURE 5**

**a-c)** RT-qPCR analysis of select transcripts in HUDEP2-Cas9 cells transduced with the indicated sgRNAs after day 5 of differentiation. Data shown are the means +/- standard deviation from n=3 biological replicates. *p* values were calculated by one way ANOVA followed by Fisher's LSD test and all comparisons are to sgNegative.

**d)** Gene overlap of RNA-seq with CUT&RUN data demonstrating odds ratios of pairwise comparisons. For both CUT&RUN and RNA-seq data, gene lists of differentially expressed or differentially bound by H3K27me3 (using nearest gene from peak center) were filtered based on FDR<0.05.

**e-g)** Gene ontology (GO) analysis of RNAseq differentially expressed genes for sgEZH2-CXC vs sgNegative (**e**), sgEZH2-SET vs sgNegative (**f**), and sgEZH2-CXC vs sgEZH2-SET **(g)**. GO terms were determined based on padj<0.05 and LFC>1.0 and ranked based on significance.

**SUPPLEMENTAL FIGURE 6**

**a-d)** RT-qPCR analysis of select transcripts in HUDEP2-Cas9 cells with Empty vector, EZH2-WT, or EZH2∆14 ectopic expression additionally transduced with negative control sgRNA or sgRNA targeting endogenous EZH2. Data shown are the means +/- standard deviation from n=3 biological replicates. *p* values were calculated by one way ANOVA followed by Fisher's LSD test.

**e**) Median fluorescence intensity (MFI) of HbF-APC measured by flow cytometry in the indicated cell lines. Bars represent mean values +/- standard deviation.

**f-h)** Validation of CUT&RUN data analysis with ChIP-qPCR. CUT&RUN tracks for H3K27me3 in the knockout rescue cell lines were compared with input normalized ChIP enrichment for n=3 biological replicates at the *SP9* (**f**), *NR2F1* (**g**), and *JPH1* (**h**) loci. Data shown are the means +/- standard deviation from n=3 biological replicates. *p* values were calculated by one way ANOVA followed by Fisher's LSD test.

**i**) CUT&RUN signal for H3K27me3 signal for the knockout rescue lines for the *LIN28B* locus.

**j)** Western blot analysis of *EZH2* knockout rescue HUDEP2 lines demonstrating that EZH2-WT can rescue H3K27me3 and LIN28B expression while EZH2Δ14 cannot.

**k)** Co-immunoprecipitation experiment in undifferentiated HUDEP2 cells expressing empty vector, 3x-FLAG-EZH2-WT, and 3xFLAG-EZH2Δ14 demonstrating that EZH2Δ14 interacts with core PRC2 subunit EED.

**l)**Co-immunoprecipitation experiment in undifferentiated HUDEP2 cells expressing empty vector, 3x-FLAG-EZH2-WT, and 3xFLAG-EZH2Δ14 demonstrating that EZH2Δ14 interacts with accessory subunits JARID2 and PHF19.

**m**) Co-immunoprecipitation experiment in HEK293T cells with ectopic expression of an empty vector, 3x-FLAG-EZH2-WT, or 3x-FLAG-EZH2Δ14 demonstrating that EZH2-WT and EZH2Δ14 can interact with core PRC2 subunits SUZ12 and RBBP4/7.

**SUPPLEMENTAL FIGURE 7**

**a)** Diagram depicting experimental workflow for targeted Cas9 gene editing in mouse fetal liver (MFL) cells carrying heterozygous humanized b-globin and Cas9 alleles. Image created in BioRender

**b)** Fraction of *HBG* transcripts (HBG/(HBB+HBG)) in MFL cells transduced with indicated sgRNAs after 48h in differentiation media. Data shown is from n=2 biological replicates

**c)** Aggregated cell counts from primary mouse FL cells from days 0-5 of expansion post spinfection with n=2 biological replicates

**d)** Flow cytometry plots of MFL transduced with the indicated sgRNAs at 24h of differentiation. Contour plots are gated FSC/SSC for viable cells and display Ter119 and CD71 with quartile populations quantified.

**e)** Inference of CRISPR edits (ICE) analysis of MFL cells transduced with sgRosa26, sgEZH2-CXC, or sgEZH2-SET sgRNAs showing successful editing by CRISPR-Cas9.

**f)** *Ezh2* full length sequencing of cDNA from fetal liver cells transduced with the indicated sgRNAs.

**SUPPLEMENTAL FIGURE 8**

**a)** Boxplots for the Fitness LFC (dropout) scores of all 100 negative control sgRNAs, sgRNAs within the EZH2-CXC domain (cut site amino acids 520-536), sgRNAs targeting the SET domain (amino acids 612-727), and all other EZH2 sgRNAs for comparison. The dashed line represents the median Fitness score for negative control sgRNAs, and the shaded area represents +/- 1 standard deviation for negative control sgRNAs. P values are calculated based on Mann-Whitney U test on shown comparisons.

**b)** Boxplots for the HbF LFC scores with the same layout as in **a**. The dashed line represents the median HbF LFC score for all negative control sgRNAs and the shaded area is +/- 1 standard deviation. P values are calculated based on Mann-Whitney U test on shown comparisons.

**c)** Two one-sided test (TOST) for the selected sgRNAs within the sgEZH2 CXC window (amino acids 520-536) comparing the Fitness LFC distribution mean +90% confidence interval (red dot and bars) to the negative control sgRNA mean (vertical dashed line). Each row bar represents an equivalence zone (+/- 0.3-1.5 LFC). Green bars represent indicate equivalence at tested interval ($p<0.05$) while grey bars indicate non-equivalence ($p>0.05$).

**d)** Two one-sided test (TOST) for the selected sgRNAs within the sgEZH2 SET window (amino acids 612-727) comparing the Fitness LFC distribution mean +90% confidence interval (blue dot and bars) to the negative control sgRNA mean (vertical dashed line). Each row bar represents an equivalence zone (+/- 0.3-1.5 LFC). All comparisons were non-significant (grey) indicating non-equivalence (p>0.05).

**e)** HUDEP2 sgRNA-GFP+ dropout assay. HUDEP2 cells were transduced with the indicated sgRNAs and the GFP+ fraction of cells was measured starting 2 days after transduction. The relative fraction of GFP+ cells was measured on the indicated days as cells were grown in HUDEP2 expansion media. Data shown are the mean fraction %GFP (normalized to %GFP at 48h post transduction for each timepoint) with error bars as standard error of mean (SEM).

**SUPPLEMENTAL FIGURE 9**

**a-c)** Differential binding analysis (DiffBind) of H3K27me3 CUT&RUN datasets using peaks called with MACS2 broadpeak. For data shown, peaks colored pink are FDR<0.05 and blue peaks are FDR>0.05.

**d-e)** Gene ontology analysis of genes associated with k-means clusters from H3K27me3 CUT&RUN data representing cluster 1 (**d**) and cluster 2 (**e**). GO processes were ranked by FDR and the top 20 GO terms enriched in each cluster are shown.

**f)** Relative GO enrichment of most significant GO terms across k-means clusters. The top 20 GO terms across all clusters are shown and the relative enrichment of each term within each cluster is shown.

**SUPPLEMENTAL FIGURE 10**

**a**) Indel tracking of CD34+ cells targeted with sgRNA-CXC at day 3, 7, and 13 of erythroid maturation. Edited PCR trace samples are shown above control PCR traces and the relative indel frequency is displayed.

**b)** Indel tracking of CD34+ cells targeted with sgRNA-SET at day 3, 7, and 13 of erythroid maturation. Edited PCR trace samples are shown above control PCR traces and the relative indel frequency is displayed.

**c)** Enucleation frequencies of CRISPR-Cas9 edited CD34+ cells following 21 day erythroid differentiation protocol. CD34+ cells were edited with the indicated sgRNAs and enucleation was measured in total CD235+ cell populations by absence of Hoechst staining.

**Supplemental Table 1:** Antibodies and key resources used in this study

**Supplemental Table 2:** MaGEcK normalized sgRNA counts from CRISPR-Cas9 screen

**Supplemental Tables 3-6:** DEseq2 output for differentially expressed genes in HUDEP2 cells

**Supplemental Tables 7-9:** DiffBind outputs for differential peak analysis of CUT&RUN data in HUDEP2 cells

**Supplemental Table 10:** sgRNAs used in this study

**Supplemental Table 11:** PCR primers used in this study

**Supplemental Tables 12-13:** Gene lists and GO terms for Cluster 1 and cluster 2 regions

### Code Availability

All code data processing were obtained from previously established pipelines which have been provided in this article.

SUPPLEMENTAL FIGURE 1

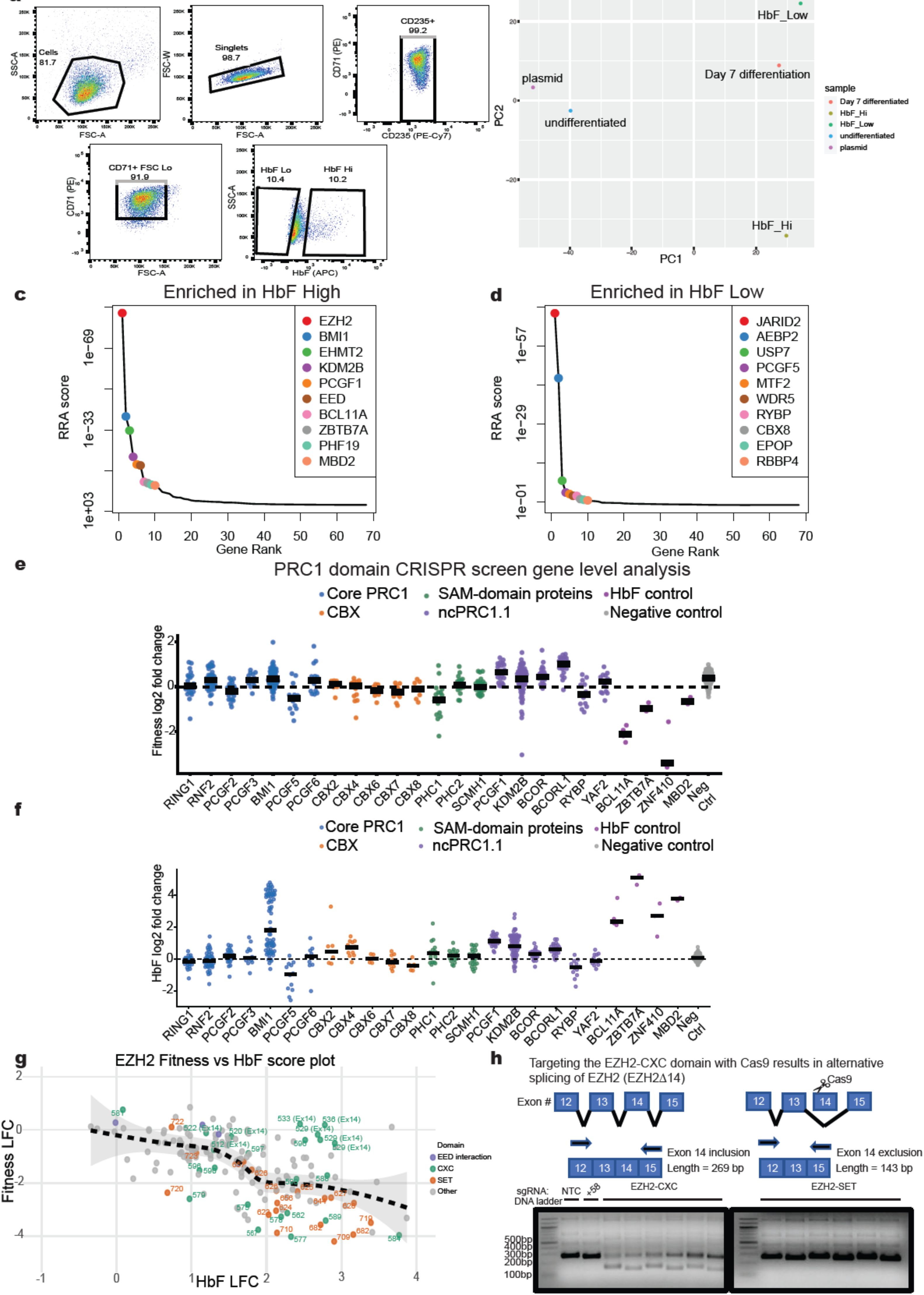

SUPPLEMENTAL FIGURE 2

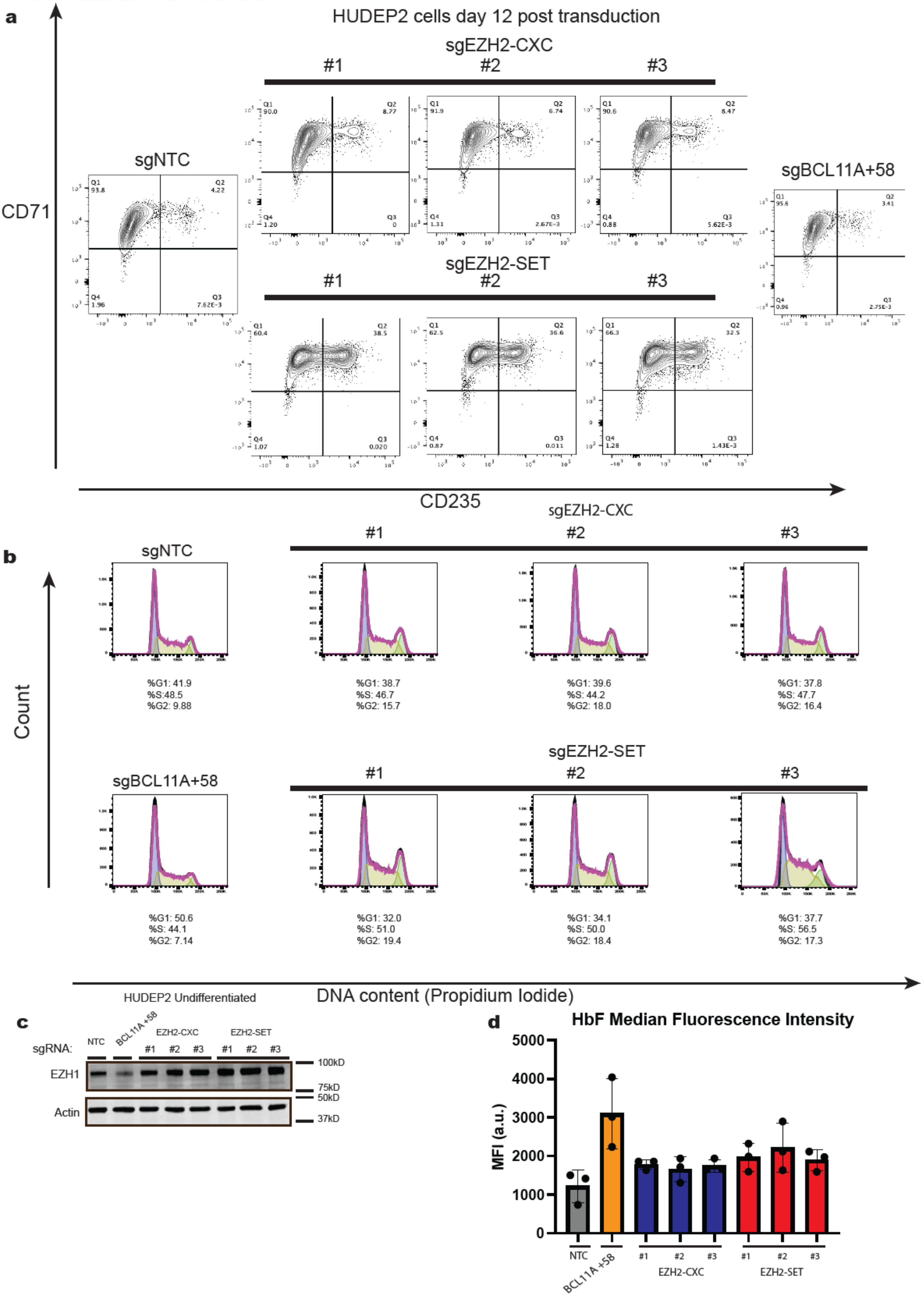

SUPPLEMENTAL FIGURE 3

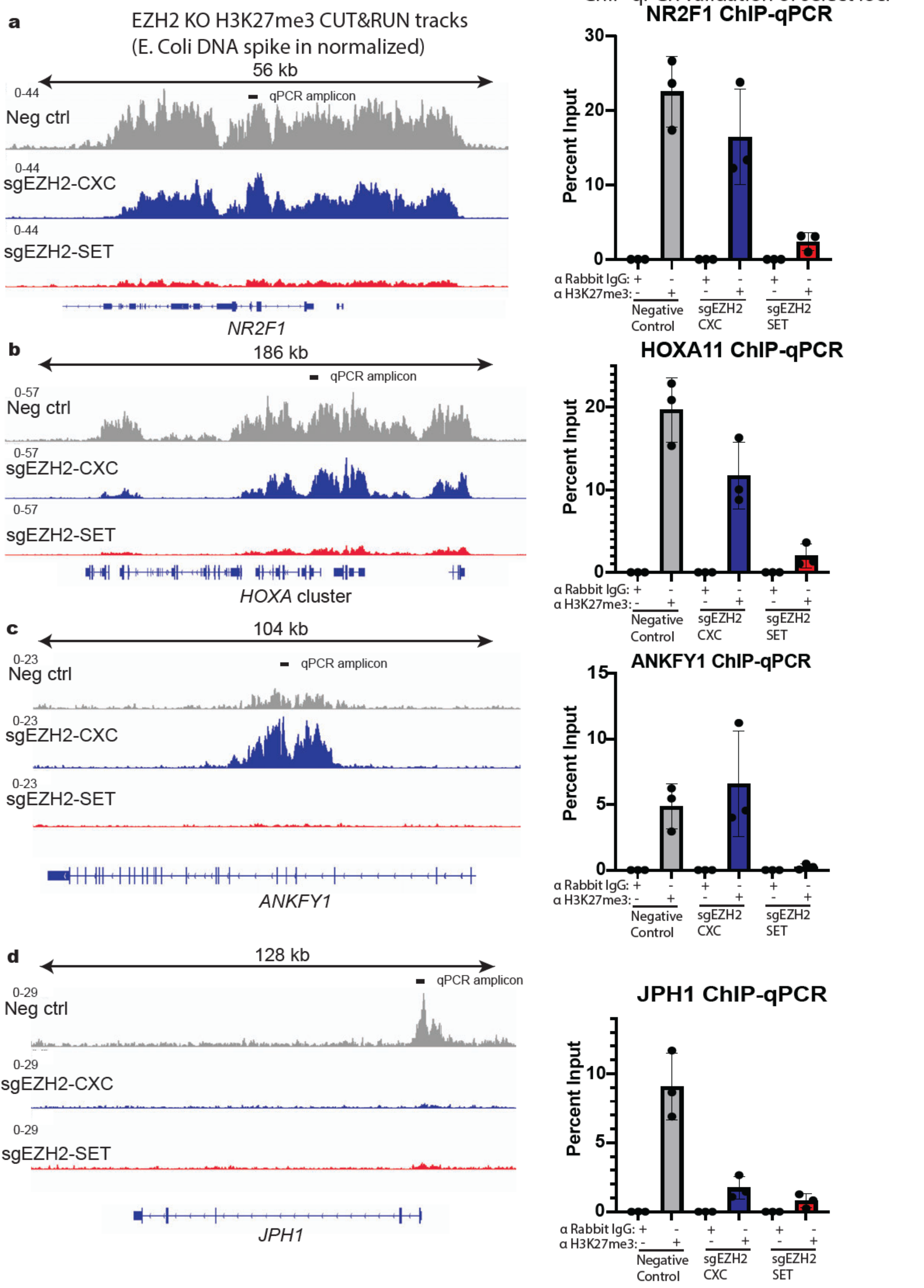

SUPPLEMENTAL FIGURE 4

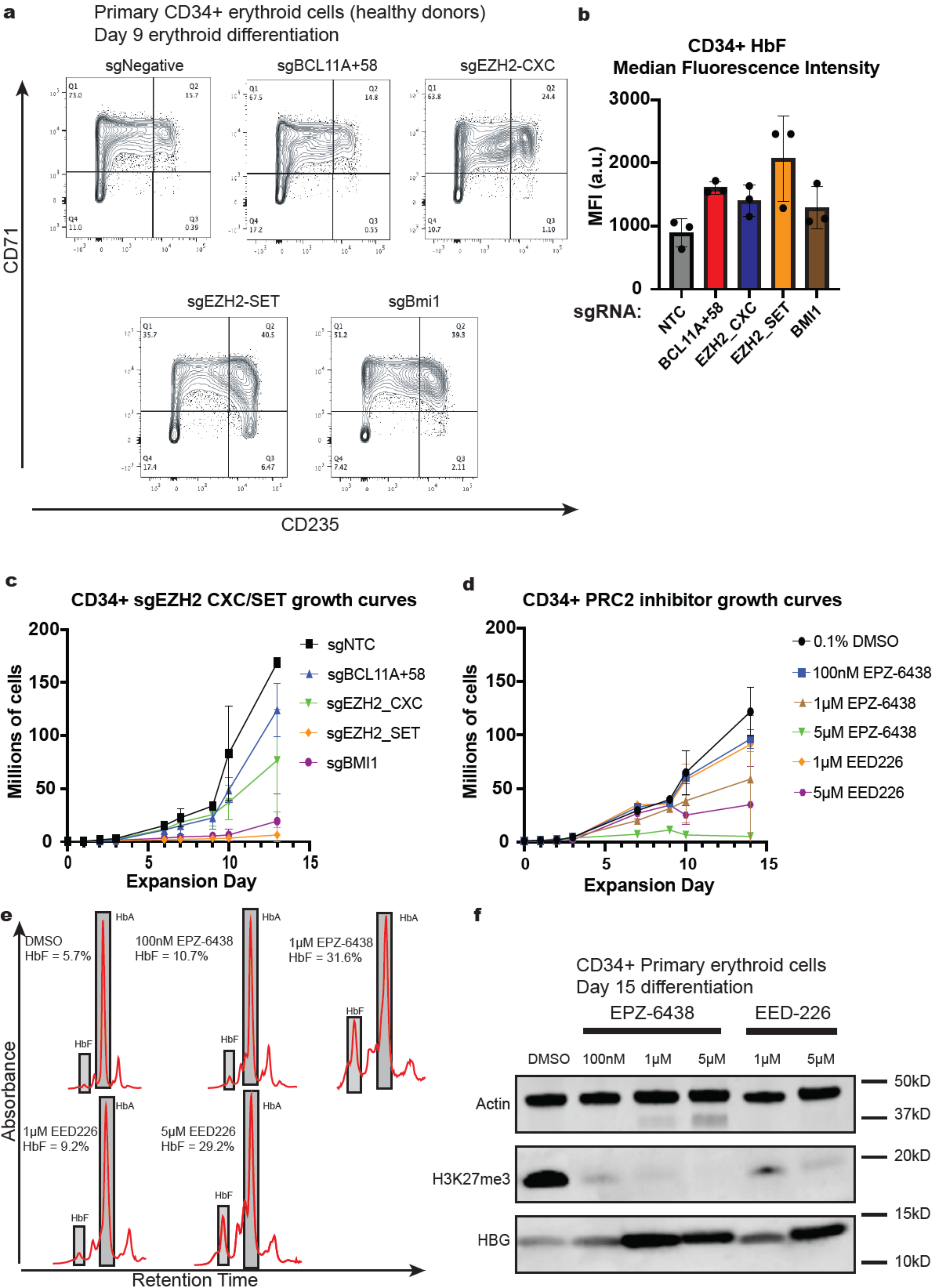

SUPPLEMENTAL FIGURE 5

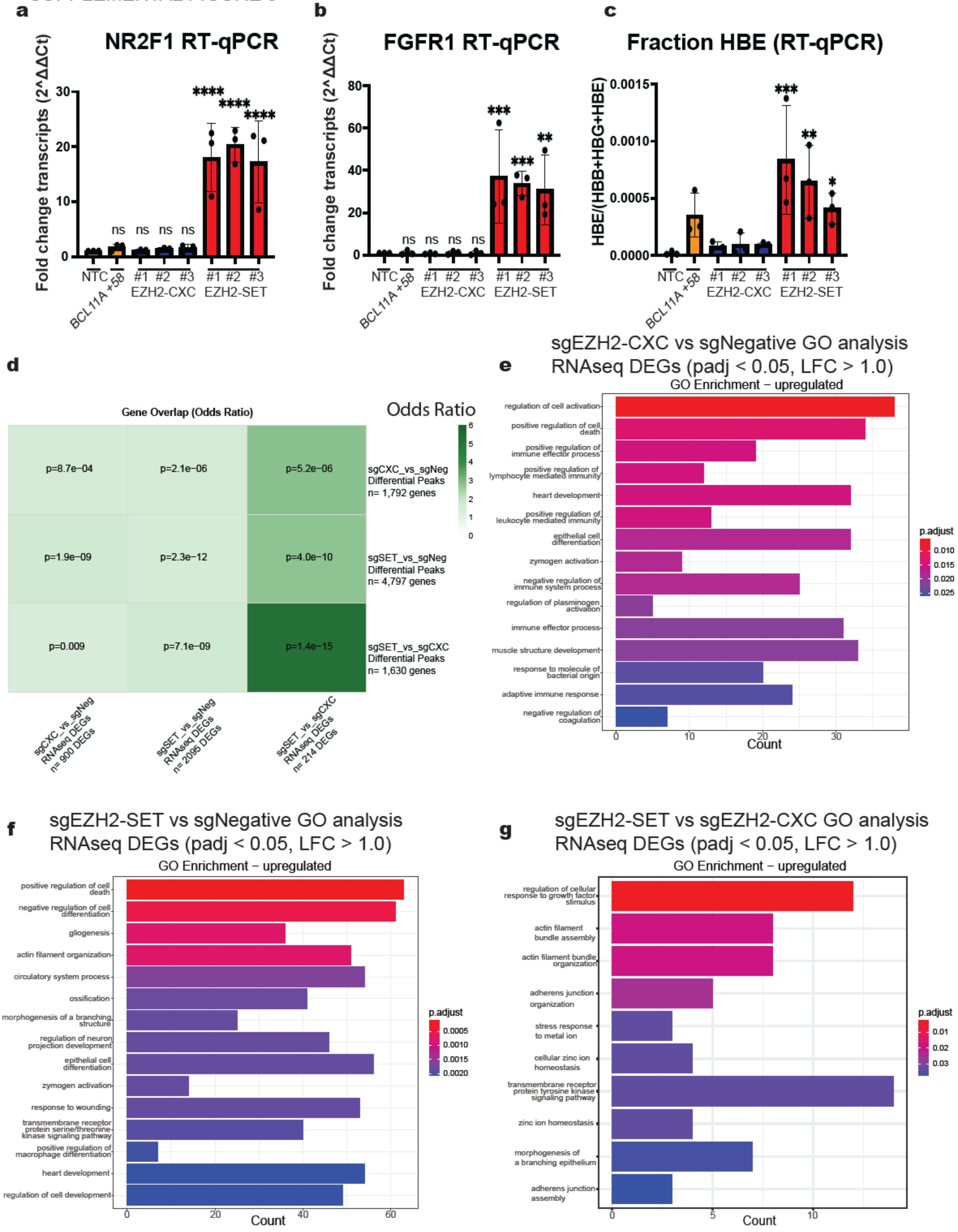

SUPPLEMENTAL FIGURE 6

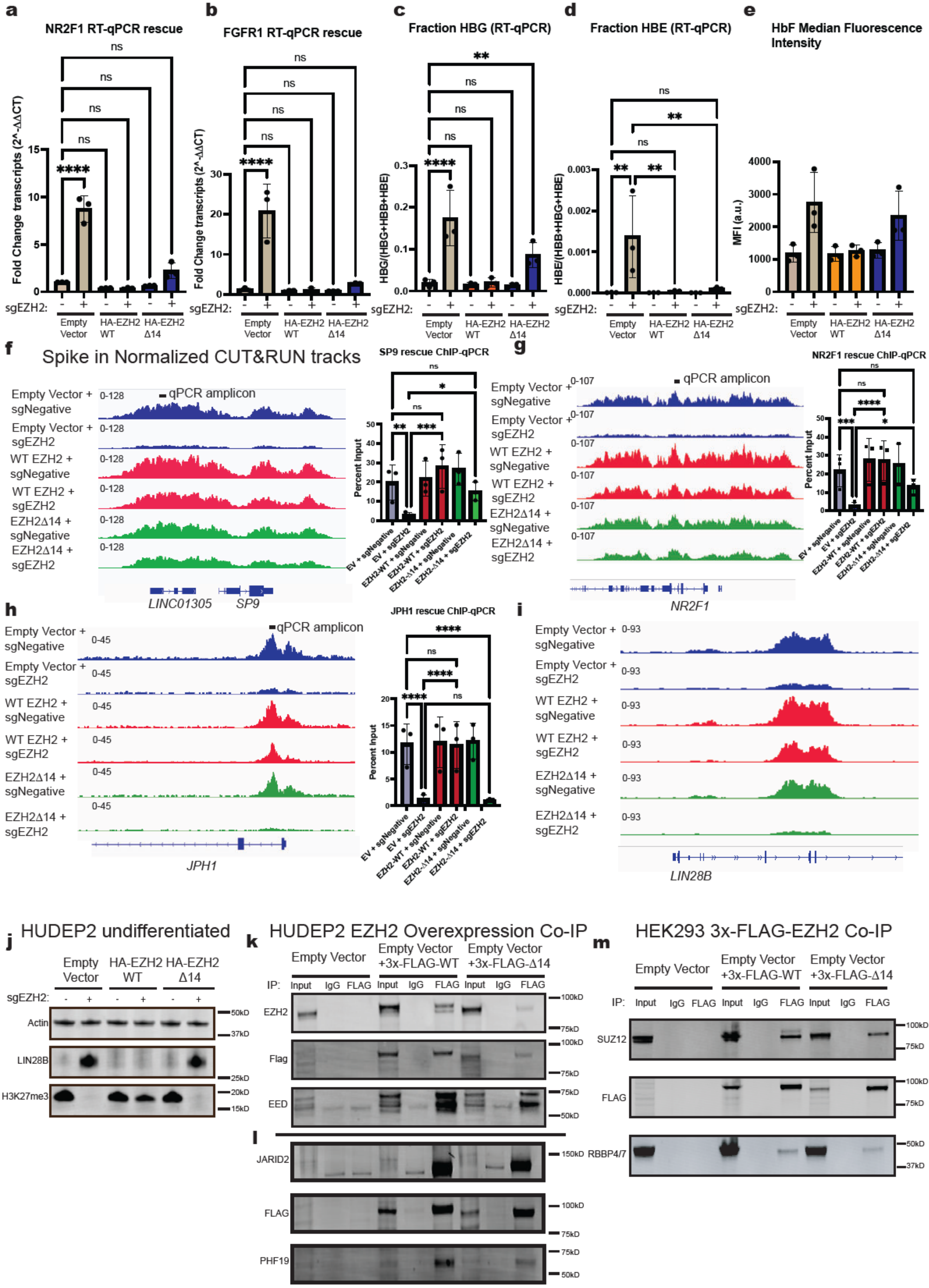

SUPPLEMENTAL FIGURE 7

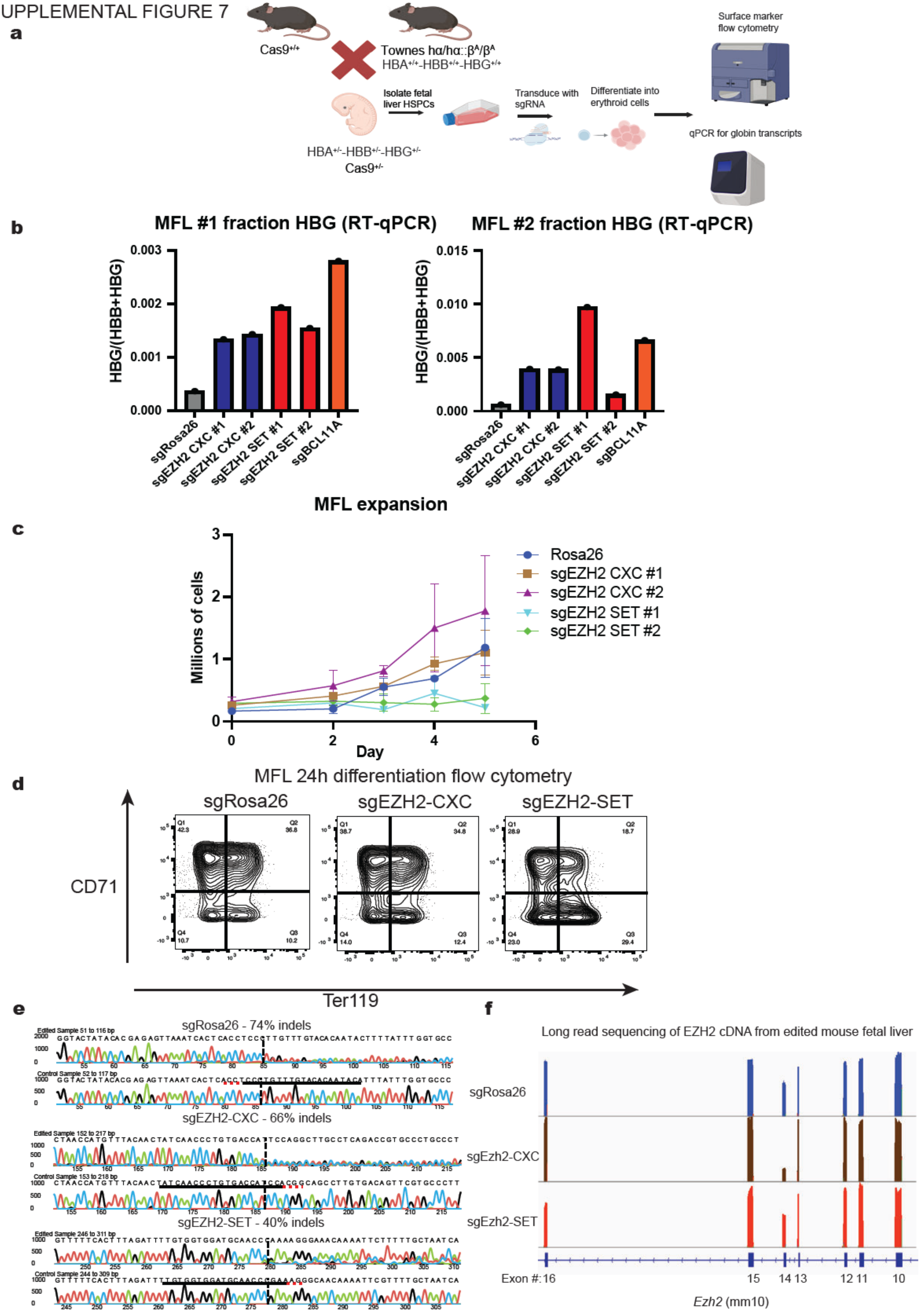

SUPPLEMENTAL FIGURE 8

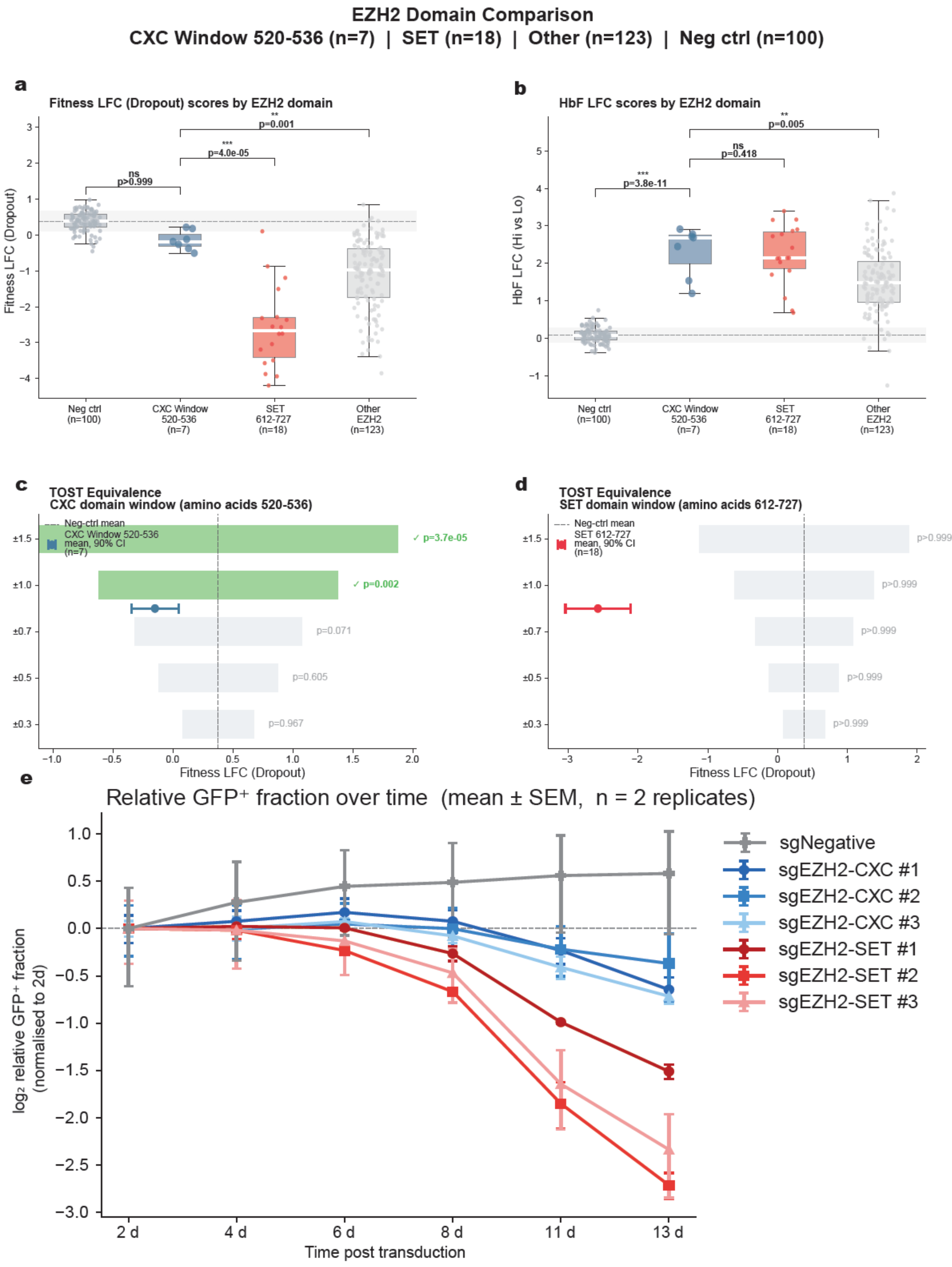

SUPPLEMENTAL FIGURE 9

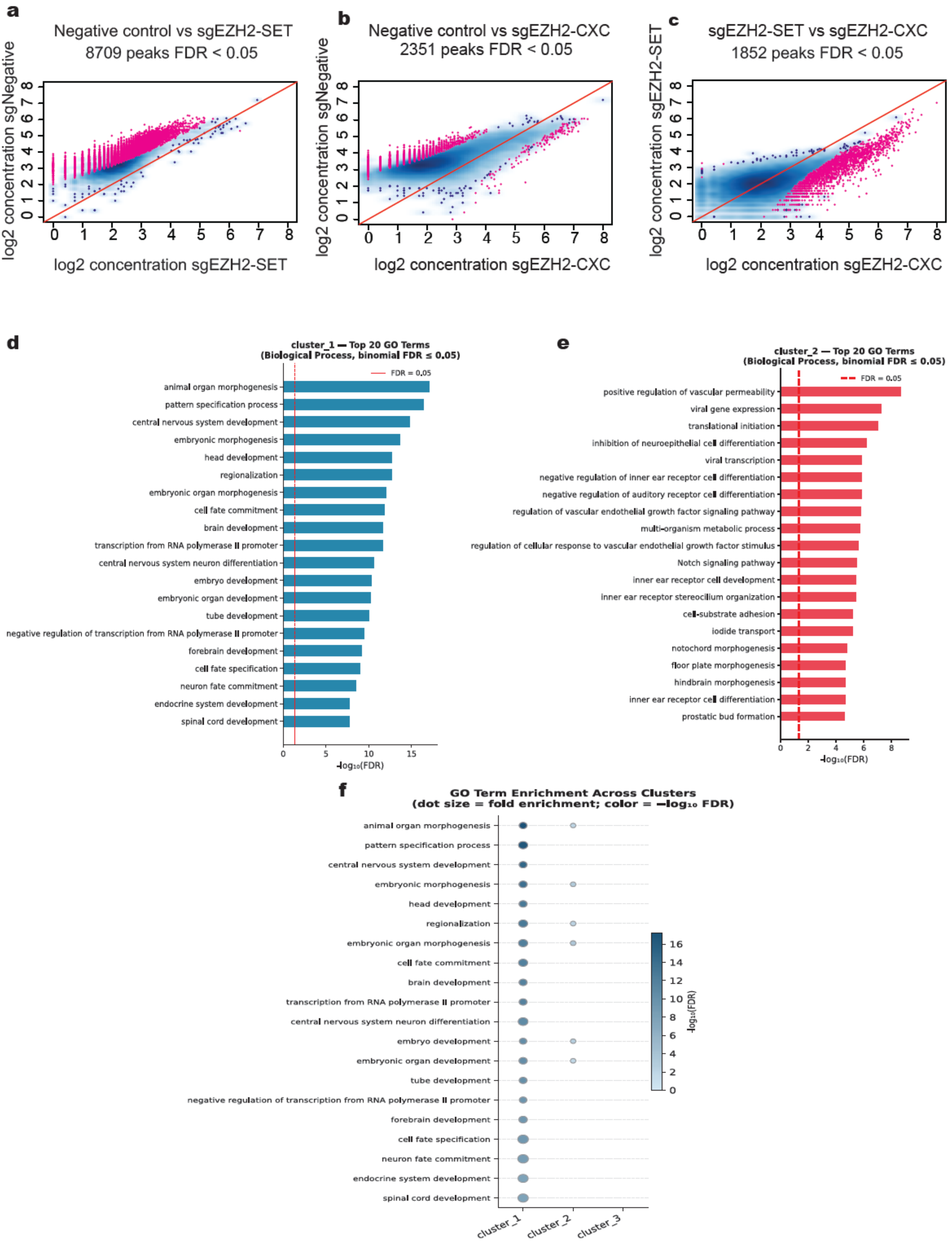

SUPPLEMENTAL FIGURE 10

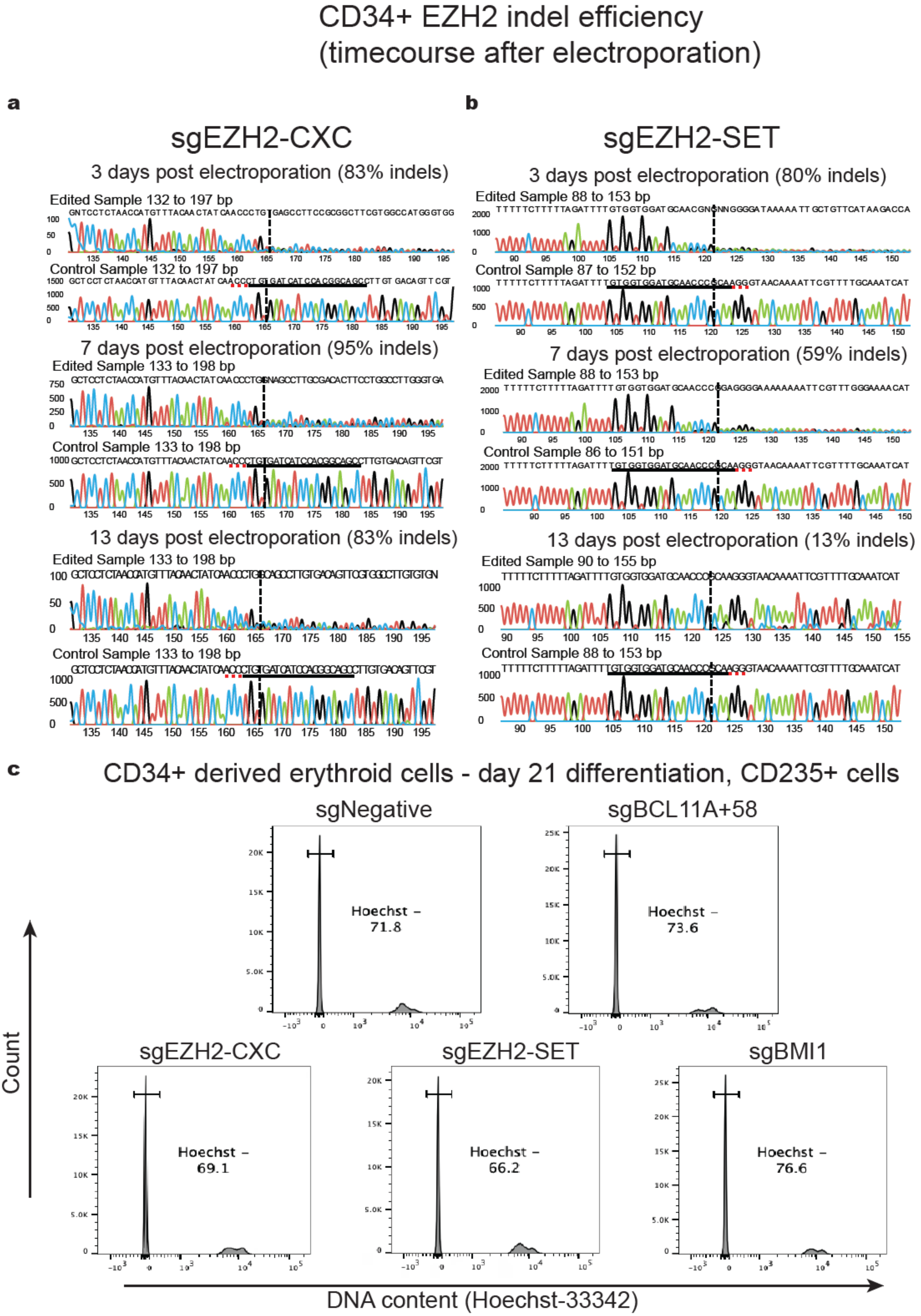
